# CAMSAP-driven microtubule release from γ-TuRC and its regulation by nucleation-promoting factors

**DOI:** 10.1101/2022.08.03.502613

**Authors:** Dipti Rai, Shasha Hua, Jooske L. Monster, Riccardo Stucchi, Kelly Stecker, Yaqian Zhang, Eugene A. Katrukha, Maarten Altelaar, Michal Wieczorek, Kai Jiang, Anna Akhmanova

## Abstract

γ-tubulin ring complex (γ-TuRC) is the major microtubule-nucleating factor. After nucleation, microtubules can be released from γ-TuRC and stabilized by other proteins, such as CAMSAPs, but the biochemical cross-talk between minus-end regulation pathways is poorly understood. Here, we reconstituted this process in vitro using purified components. We found that all CAMSAP proteins could bind to the minus-ends of γ-TuRC-attached microtubules. CAMSAP2 and CAMSAP3, which decorate and stabilize growing minus ends, but not the minus-end tracking protein CAMSAP1 induced microtubule release from γ-TuRC. CDK5RAP2, a γ-TuRC-interactor, and CLASP2, a regulator of microtubule growth, stimulated γ-TuRC-dependent microtubule nucleation, but only CDK5RAP2 inhibited CAMSAP-driven microtubule detachment by suppressing CAMSAP binding to γ-TuRC-anchored minus ends. CDK5RAP2 also improved γ-TuRC selectivity for 13-rather than 14-protofilament microtubules in microtubule-capping assays. Our results support a model whereby CAMSAPs exploit an imperfect attachment between γ-TuRC and the nucleated microtubule to promote minus-end elongation and release, whereas CDK5RAP2 improves the fit between γ-TuRC and 13-protofilament microtubules and enhances nucleation.

## Introduction

Microtubule organization in animal cells is a major determinant of cell architecture and polarity^1^. This organization critically depends on the activity of microtubule-organizing centers (MTOCs) – structures that can nucleate microtubules and stabilize and anchor their minus ends^2–4^. The major microtubule-nucleating factor in cells is the γ-tubulin ring complex (γ-TuRC) – a cone-shaped macromolecular assembly that is required for the kinetically dominant pathway of generating new microtubules from tubulin dimers^5, 6^. γ-TuRC localization and activity are controlled by a variety of tethering and nucleation-promoting factors, such as augmin, pericentrin, CDK5RAP2 and chTOG^6, 7^. γ-TuRC can also cap microtubule minus ends^8^ and participate in their anchoring, possibly with the aid of additional MTOC components^9^. An alternative well-studied pathway of minus-end stabilization and anchoring depends on the members of CAMSAP/Patronin family^10–13^. These proteins specifically recognize free, uncapped microtubule minus ends because their signature domain, CKK, binds to a minus-end-specific site between flared protofilaments^14^. Recently, a role of CAMSAP2 in γ-tubulin-independent microtubule nucleation has also been proposed^15^.

Importantly, since in γ-TuRC-capped microtubules protofilaments at the minus ends are straight^16, 17^, they should not be able to bind to CAMSAPs. However, recent structural work revealed that purified γ-TuRCs are strikingly asymmetric, and their structure does not fully match with that of a complete 13-protofilament microtubule^18–21^. This finding raises the possibility that γ-TuRC-nucleated microtubules may not be fully attached to their template, and some protofilaments might have a flared conformation that would permit CAMSAP binding. Furthermore, since the microtubule-nucleating activity of purified γ-TuRC turned out to be quite low^18, 22^, a potential mechanism of stimulating nucleation would be to alter γ-TuRC conformation to make it more similar to the microtubule structure^6, 7, 23, 24^. To explore these possibilities, we have set up in vitro reconstitution assays and confirmed that purified γ-TuRC has by itself a rather low activity that can be enhanced by nucleation-promoting factors, microtubule polymerase chTOG, and γ-TuRC associated protein CDK5RAP2^25–27^. We further established that the microtubule-nucleation activity of γ-TuRC can also be stimulated by CLASP2, a protein known to strongly enhance microtubule outgrowth from stabilized microtubule seeds^28^.

We then set up assays where the activities of γ-TuRC, CAMSAPs and CDK5RAP2 or CLASP2 could be observed simultaneously. We found that while microtubules almost never detached from γ-TuRC when it was present alone or together with CDK5RAP2 or CLASP2, CAMSAPs could bind to a subset of γ-TuRC-anchored minus ends and trigger their detachment. While minus-end binding was observed with all three CAMSAPs, microtubule release from γ-TuRC was only induced by CAMSAP2 and 3, which decorate and stabilize growing microtubule minus ends, but not by CAMSAP1, which tracks growing minus ends without decorating them^29^. We also found that CDK5RAP2, but not CLASP2 suppressed CAMSAP binding and subsequent microtubule release. We reasoned that this could be due to the ability of CDK5RAP2 to alter γ-TuRC conformation so that it would fit the regular microtubule structure better. We found support for this idea in microtubule-capping assays, where CDK5RAP2 made γ-TuRC more selective for 13-protofilament microtubules. Together, our results indicate that CAMSAPs can bind to a subpopulation of microtubule minus ends anchored at γ-TuRC, likely because they are only partially attached to their template. By decorating and stabilizing the growing microtubule minus ends, CAMSAP2 and 3 help to generate the force required to push the γ-TuRC cap away. This process can be counteracted by the γ-TuRC-binding factor CDK5RAP2, possibly because it improves the fit between the γ-TuRC and a microtubule.

## Results

### Human CDK5RAP2, CLASP2 and chTOG promote microtubule nucleation by purified γ-TuRC

To obtain purified γ-TuRC, we have used CRISPR-Cas9-mediated gene editing to generate a homozygous knock-in HEK293T cell line where the GCP3-encoding gene was modified by a C-terminal insertion of the GFP and a Twin-Strep-tag (two Strep-tag II in tandem, abbreviated as GFP-SII, Extended data Fig. 1a,b). Western blotting of this cell line showed that the whole GCP3 pool was shifted up by ∼30 kDa (Extended data Fig. 1c), and fluorescence microscopy revealed GFP signal was present diffusely in the cytoplasm and concentrated at the centrosome, as expected for a γ-TuRC component (Extended data Fig. 1d). Twin-Strep-tag based purification (Extended data Fig. 1e) yielded protein complexes, which, based on Western blotting and mass spectrometry, contained all expected γ-TuRC components (Extended data Fig. 1f, g and Supplementary Table 1). In these γ-TuRC preparations, we detected two proteins known to co-purify with γ-TuRC, NEDD1 (Neural precursor cell expressed developmentally down-regulated protein 1) and NME7 (Nucleoside diphosphate kinase 7) ^18–20, 26, 30, 31^ (Extended data Fig. 1g), but no other known γ-TuRC binding partners or microtubule nucleation-promoting factors. 3D reconstruction of negative stain transmission EM micrographs of purified γ-TuRCs showed a cone-like complex (Fig. 1a and see below), matching the recently published γ-TuRC structures ^17–21^. We also characterized the fluorescence intensity of purified γ-TuRC and compared it to that of single GFP molecules and GFP-EB3, which is known to be a dimer ^32^. In these measurements, GFP-EB3 was ∼1.7x brighter than GFP, while GCP3-GFP-containing γ-TuRC showed a rather broad distribution and was on average ∼4.1x brighter than GFP (Fig. 1b). This fits with the fact that γ-TuRCs derived from cells where all GCP3 subunits bear a GFP tag would be expected to contain five GCP3-GFP subunits ^18–21, 24^.

**Fig. 1.**
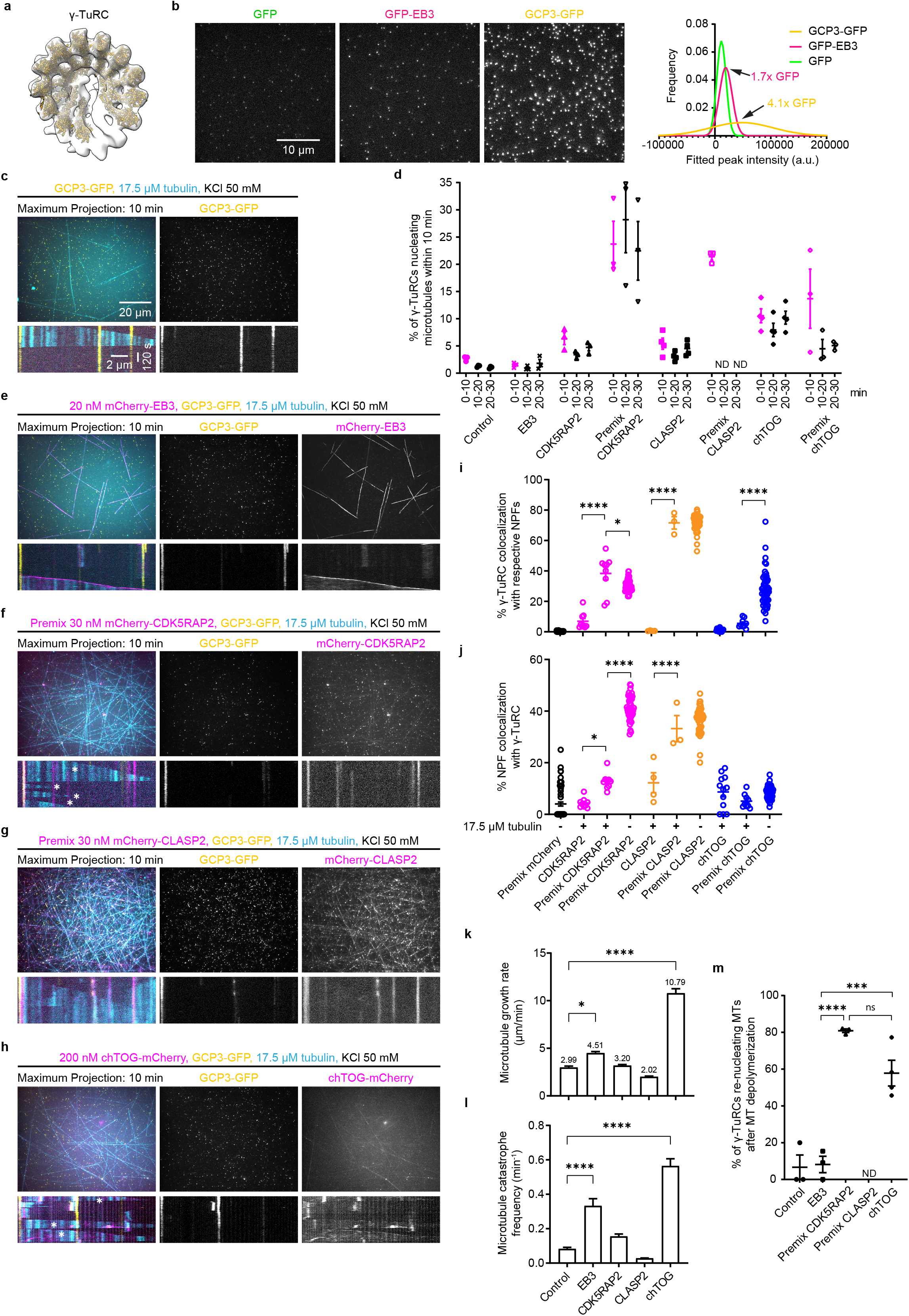
Human CDK5RAP2, CLASP2 and chTOG promote microtubule nucleation by purified γ-TuRC. **a,** A 3D reconstruction of the γ-TuRC from negative-stain EM data and rigid body fit of repeating γ-tubulin/GCP2 subcomplexes (from PDB ID: 6V6S ^20^) individually docked into the γ-TuRC density map. Fits for two subcomplexes at the γ-TuRC “seam” were not reliable and are therefore omitted for clarity. 3D reconstruction was generated from 12,851 particles (Also see **Extended data Fig. 4i** for details). **b,** Left: representative images of single molecules of the indicated purified proteins (GFP, GFP-EB3, GCP3-GFP) attached to coverslips. Right: corresponding non-linear regression fits of histograms of single molecule fluorescence intensities using least squares method. Data are pooled from three independent experiments: n=21,349 for GFP (green), n=28,776 for GFP-EB3 (magenta) and n=24,216 for GCP3-GFP (yellow), where n is the number of molecules analyzed. **c, e-h,** Top: maximum intensity projections of 10 min time-lapse videos, acquired after 20 min of incubation, showing microtubules (cyan) nucleated from γ-TuRC (GCP3-GFP, yellow) in the presence of either 17.5 μM tubulin (17 μM unlabeled porcine tubulin and 0.5 μM HiLyte647-tubulin) and 50 mM KCl only (**c**), or together with the indicated proteins shown in magenta (**e-h**). For **f** and **g**, γ-TuRC was also preincubated with the same concentration of indicated proteins before immobilization on the coverslips. Bottom: representative kymographs illustrating dynamics of microtubules nucleated from γ-TuRC and microtubule re-nucleation events (asterisks) from experiments shown in top panels. **d**, Quantification of average microtubule nucleation efficiency of γ-TuRC in the presence of either 17.5 μM tubulin alone (n=3), or together with 20 nM mCherry-EB3 (n=3); 30 nM mCherry-CDK5RAP2 (n=3); 30 nM mCherry-CDK5RAP2, also preincubated with γ-TuRC (n=3); 30 nM mCherry-CLASP2 (n=4); 30 nM mCherry-CLASP2, also preincubated with γ-TuRC (n=3); 200 nM chTOG-mCherry (n=4); 200 nM chTOG-mCherry, also preincubated with γ-TuRC (n=3), where n is the number of independent experiments. Representative images are shown in **c,e-h** (top panels), and **Extended data Fig. 2c,e**. The plot presents mean ± s.e.m., and each data point represents a single field of view from which % of γ-TuRCs nucleating microtubules were quantified for the given time point from an individual experiment. Data points in magenta (0-10 min) were acquired from a smaller field of view and are not directly comparable to the data points in black (10-20 min and 20-30 min) acquired from a bigger field of view shown in top panels of **c, e-h**. **i,j,** Quantification of colocalization of γ-TuRC with indicated nucleation-promoting factors (NPFs)(**i**) or colocalization of indicated NPFs with γ-TuRC (**j**) under experimental conditions shown in **c, e-h,** and **Extended data Fig. 2d.** The plots show data for γ-TuRC premixed with mCherry without tubulin (n=63, N=3); γ-TuRC with mCherry-CDK5RAP2 and 17.5 μM tubulin (n=9, N=3); γ-TuRC premixed with mCherry-CDK5RAP2 with (n=9, N=3) and without 17.5 μM tubulin (n=47, N=3); γ-TuRC with mCherry-CLASP2 and 17.5 μM tubulin (n=4, N=4); γ-TuRC premixed with mCherry-CLASP2 with (n=3, N=3) and without 17.5 μM tubulin (n=48, N=3); γ-TuRC with chTOG-mCherry and 17.5 μM tubulin (n=12, N=4); γ-TuRC premixed with chTOG-mCherry with (n=9, N=3) and without 17.5 μM tubulin (n=51, N=3), where n is the number of fields of view analyzed and N is the number of independent experiments. The plots present mean ± s.e.m., and individual data points represent % of γ-TuRC molecules colocalizing with the indicated protein or vice versa in a single field of view. One-way ANOVA with Tukey’s multiple comparisons test was used to compare the means with each other (**i:** *p=0.0199; ****p< 0.0001, **j:** *p=0.0246; ****p< 0.0001). **k,l,** Quantification of average plus-end growth rate (**k**) and catastrophe frequency (**l**) of microtubules nucleated from γ-TuRC in the presence of either 17.5 μM tubulin alone (n=25), or together with mCherry-EB3 (n=36); mCherry-CDK5RAP2 (n=52); mCherry-CLASP2 (n=60); chTOG-mCherry (n=78), where n is the number of growth events analyzed. The bar plots present mean ± s.e.m. from three independent experiments. Representative kymographs are shown in **c,e,h** (bottom panels) and **Extended data Fig. 2c,e**. One-way ANOVA with Dunnett’s multiple comparisons test was used to compare the means with control (**k:** *p=0.0444; ****p< 0.0001, **l:** ****p< 0.0001). **m,** Quantification of microtubule re-nucleation efficiency of γ-TuRCs analyzed from experiments shown in **c, e-h.** The plot shows data for γ-TuRC with 17.5 μM tubulin only (n=3, m=10); together with mCherry-EB3 (n=3, m=26); premixed mCherry-CDK5RAP2 (n=3, m=107); premixed mCherry-CLASP2 (n=3, m=0) and chTOG-mCherry (n=4, m=178), where n is the number of independent experiments analyzed and m is the number of γ-TuRCs which nucleated microtubules that underwent depolymerization, pooled from three 10 min videos of a single experiment. The plot presents mean ± s.e.m., and individual data points represent % of γ-TuRCs re-nucleating microtubules in a single experiment. One-way ANOVA with Tukey’s multiple comparisons test was used to compare the means with each other (ns, not significant, p=0.0704; ***p=0.0007; ****p< 0.0001). ND, could not be determined.

We next immobilized purified GFP-tagged γ-TuRCs on coverslips using a biotinylated GFP nanobody, observed microtubule nucleation in the presence of Rhodamine-labeled tubulin by Total Internal Reflection Fluorescence (TIRF) microscopy (Extended data Fig. 1h) and counted the percentage of γ-TuRCs nucleating microtubules within three consecutive 10 min periods. In the presence of 17.5 µM tubulin, only ∼1-3% of γ-TuRC could nucleate microtubules per 10 minutes of observation (Fig. 1c,d), which is similar to previous studies using purified γ-TuRCs obtained from other cell types using different tagging and purification approaches^18, 22^. We then set out to investigate the impact of several microtubule- or γ-TuRC-binding proteins on γ-TuRC-mediated microtubule nucleation (Fig. 1d-j). The addition of mCherry-EB3 did not affect the nucleation efficiency although it increased both microtubule growth rate and catastrophe frequency (Fig. 1d,e,k,l and Extended data Fig. 2a). In contrast, three nucleation-promoting factors, CDK5RAP2, chTOG and CLASP2, could potentiate microtubule nucleation, both when added to γ-TuRC immobilized on coverslips or when additionally preincubated with γ-TuRC in solution (“Premix”) before immobilization (Fig. 1d, f-h, Extended data Fig. 2b,c,e,f, and see Supplementary Tables 2-4 for mass spectrometry-based characterization of the purified proteins).

Full-length mCherry-CDK5RAP2 increased microtubule nucleation ∼3 fold when added to immobilized γ-TuRCs and more than 20-fold (up to ∼35% nucleation efficiency) when additionally preincubated with γ-TuRC (Fig. 1d,f and Extended data Fig. 2b,c). This was likely because preincubation greatly increased the percentage of γ-TuRCs colocalizing with CDK5RAP2 (4-6 fold), which in preincubated samples could reach 30-40%, both in the presence and absence of free tubulin (Fig. 1f,i,j and Extended data Fig. 2c,d). These data confirm that CDK5RAP2 directly interacts with γ-TuRC ^26^ and suggest that in our assays, it can activate the majority of γ-TuRCs to which it binds. The percentage of CDK5RAP2-positive dots colocalizing with γ-TuRCs was rather low in these experiments (∼13% with tubulin, ∼41% without tubulin), likely because CDK5RAP2 was present in excess, or due to the autoinhibitory regulation of CDK5RAP2 that is controlled by its phosphorylation state^33, 34^. Compared to the samples with tubulin alone or with mCherry-EB3, mCherry-CDK5RAP2 had no significant effect on microtubule growth rate or catastrophe frequency, but strongly increased the frequency of microtubule re-nucleation after depolymerization, indicating that CDK5RAP2 can maintain γ-TuRC in an active state (Fig. 1k-m).

We also observed a strong increase in microtubule nucleation efficiency with mCherry-CLASP2 (Fig. 1d,g and Extended data Fig. 2b), a protein previously shown to promote microtubule outgrowth from stabilized seeds^28^, but never tested for interaction with γ-TuRC. The activating effect of mCherry-CLASP2 was similar in magnitude to that of CDK5RAP2 and was again stronger after preincubation (Fig. 1d,g and Extended data Fig. 2e). mCherry-CLASP2 also strongly colocalized with γ-TuRC, even in the absence of free tubulin, suggesting that it might bind to γ-TuRC directly (Fig. 1i,j and Extended data Fig. 2d). It should be noted that since CLASP2 strongly suppresses catastrophes, and therefore microtubules become very long^28^ (Fig. 1g,l), it was not possible to examine microtubule re-nucleation from the same γ-TuRC or analyze premixed CLASP2-γ-TuRC samples that were incubated longer than 10 minutes due to very high microtubule density.

Finally, we also examined the effect of chTOG, because it can weakly promote microtubule nucleation from free tubulin^35^ and strongly promote γ-TuRC-dependent microtubule nucleation^18^, and its *Xenopus* homolog XMAP215 can synergize with γ-TuRC^25^ and promote outgrowth from seeds^36^. We confirmed that chTOG enhanced γ-TuRC-dependent microtubule nucleation, although unlike CDK5RAP2 and CLASP2, the effect was similar with and without preincubation (Fig. 1d,h and Extended data Fig. 2b) and not as strong as previously published^18^. This could be due to the differences in experimental conditions, but was unlikely to be due to the low activity of chTOG, as it strongly increased the growth rate and catastrophe frequency in our assays (Fig. 1k,l), in line with the published data ^37, 38^. Colocalization of chTOG-positive dots with γ-TuRC was lower than that of CDK5RAP2 or CLASP2, but increased in the absence of free tubulin (Fig. 1i,j), with which chTOG interacts through its TOG domains^39^. It should be noted that when chTOG was preincubated with γ-TuRC, it formed clusters that sequestered γ-TuRC (Extended data Fig. 2d), and therefore we did not premix chTOG and γ-TuRC in the experiments described from here onwards. Interestingly, similar to CDK5RAP2, chTOG potentiated repeated microtubule nucleation from the same γ-TuRC (Fig. 1m).

For all these proteins, we also tested whether increasing their concentration in conditions without premixing would boost nucleation efficiency but found this not to be the case (Extended data Fig. 2a,c,e,f). This indicates that in the conditions tested, nucleation is limited by the activity and/or surface interactions of γ-TuRCs rather than the availability of nucleation-promoting factors. We conclude that purified γ-TuRC can nucleate microtubules in vitro in a manner dependent on various interactors.

### CAMSAP3 triggers microtubule release from γ-TuRC

In the assays described above, microtubule minus ends did not grow because they were stably capped and almost never released from γ-TuRCs that nucleated them, though the frequency of release events was slightly increased by chTOG (from ∼1 to∼2.5-3%) (Extended data Fig. 3a-c). We then set out to examine whether γ-TuRC prevents CAMSAPs from binding to microtubule minus ends, initially by using CAMSAP3, which among the CAMSAP family members has been suggested to have the highest minus-end affinity in vitro^29^. To recapitulate in vitro CAMSAP3 specificity for growing microtubule minus ends, the ionic strength of the buffer needs to be sufficiently high (MRB80 buffer supplemented with 80 mM KCl, as opposed to 50 mM KCl used above). Since high ionic strength suppresses microtubule assembly^40^, we increased tubulin concentration to 25 µM and obtained ∼1.5-3% nucleation efficiency with and without SNAP-AF647-CAMSAP3 (Fig. 2a,b and Extended data Fig. 3d). Also in these buffer conditions, very little microtubule release from γ-TuRC was detected in the absence of CAMSAP3 (Fig. 2c,d). Strikingly, when SNAP-AF647-CAMSAP3 was added, microtubule nucleation events were frequently followed by specific binding of SNAP-AF647-CAMSAP3 to the γ-TuRC-associated microtubule minus end and its subsequent elongation (Fig. 2e,f). In some cases, the microtubule remained attached to the glass surface in the vicinity of γ-TuRC that nucleated it (e.g. Fig. 2e and Supplementary Video 1), while in other cases the minus end detached from the surface and the microtubule floated away from the γ-TuRC (Fig. 2f and Supplementary Video 1). Approximately 25% of all γ-TuRC-nucleated microtubules acquired a CAMSAP3 signal at their minus end (Fig. 2g and Extended data Fig. 3e), and approximately half of these microtubules initiated minus-end growth (Fig. 2h). As a result, the percentage of microtubules released from γ-TuRC increased more than 10-fold, to 15% (Fig. 2d). The time interval between the initial CAMSAP3 binding and the onset of microtubule minus end elongation varied between 50-350s (Fig. 2i). After microtubule release, the same γ-TuRC could sometimes initiate growth of a new microtubule (Extended data Fig. 3f).

**Fig. 2.**
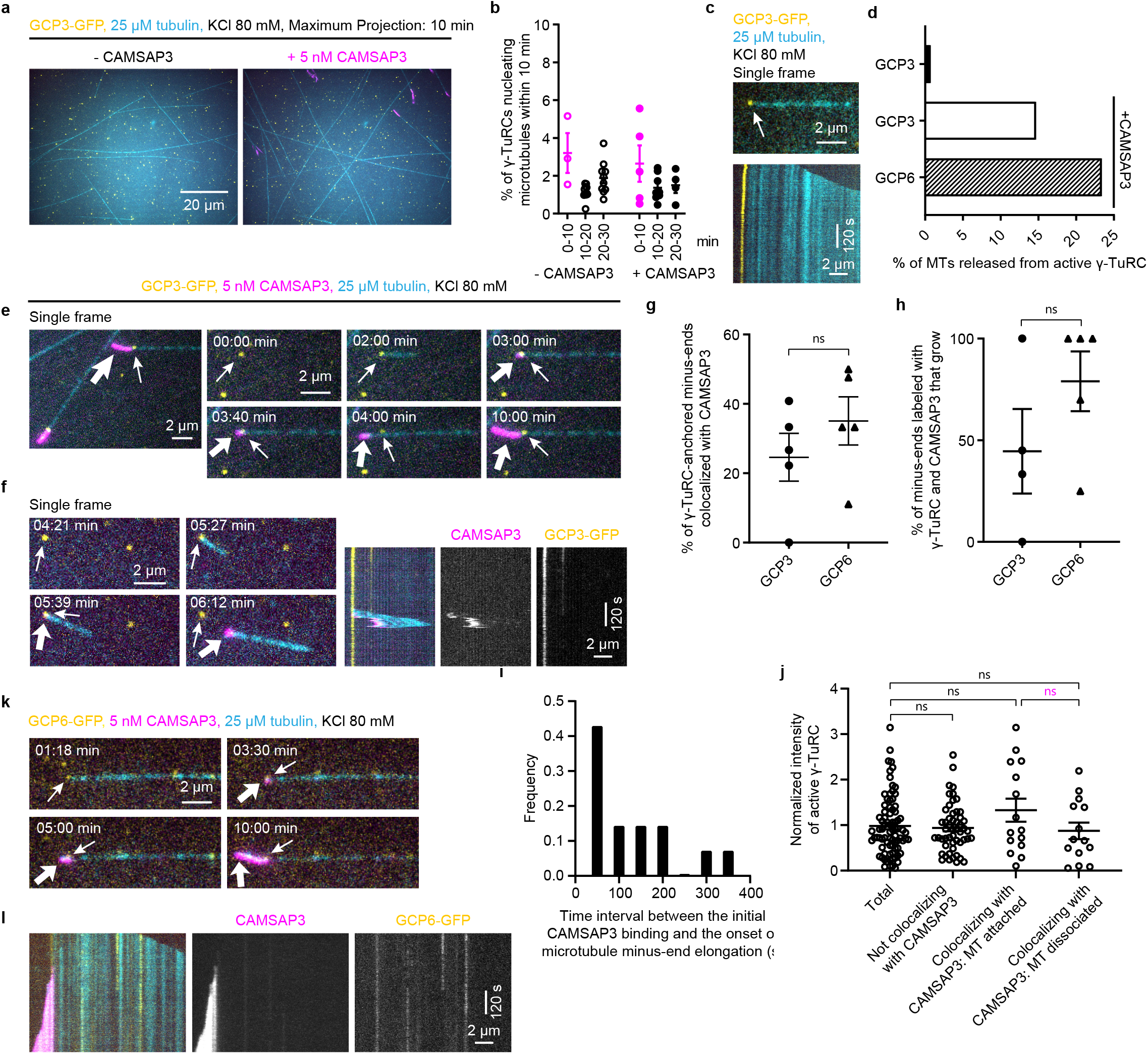
CAMSAP3 triggers microtubule release from γ-TuRC. **a,** Maximum intensity projections of 10 min time-lapse videos, acquired after 10 min of incubation, showing microtubules (cyan) nucleated from γ-TuRC (GCP3-GFP, yellow) in the presence of either 25 μM tubulin (24.5 μM unlabeled porcine tubulin and 0.5 μM rhodamine-tubulin) and 80 mM KCl only (left), or together with 5 nM SNAP-AF647-CAMSAP3 (magenta) (right). **b,** Quantification of average microtubule nucleation efficiency of γ-TuRC in the presence of either 25 μM tubulin alone (for 0-10 min, n=3; for 10-20 min, n=10; for 20-30 min, n=10, analyzed from three independent experiments) or together with SNAP-AF647-CAMSAP3 (for 0-10 min, n=5; for 10-20 min, n=8; for 20-30 min, n=4, analyzed from five independent experiments), where n is the number of fields of view analyzed. Representative images are shown in **a**. The plot presents mean ± s.e.m., and each data point represents a single field of view from which % of γ-TuRCs nucleating microtubules were quantified for the given time point from an individual experiment. Data points in magenta (0-10 min) were acquired from a smaller field of view and are not directly comparable to the data points in black (10-20 min and 20-30 min) acquired from a bigger field of view shown in **a**. **c,** Single frame (top, arrow points to a γ-TuRC-anchored microtubule) and representative kymograph (bottom) from the experiment shown in **a** without CAMSAP3. **d,** Plot showing frequency of microtubule release from active γ-TuRC in the absence (n=136 for γ-TuRC purified using GCP3-GFP-SII, pooled from three independent experiments) or presence of SNAP-AF647-CAMSAP3 (n=102 for γ-TuRC purified using GCP3-GFP-SII, pooled from five independent experiments; and n=47 for γ-TuRC purified using GCP6-GFP-SII, pooled from five independent experiments), where n is the number of active γ-TuRCs analyzed. Representative images are shown in **c,e,k**. **e,** Single frame and cropped images, at indicated time points in min, from a 10 min time-lapse video showing microtubule (cyan) nucleation from γ-TuRC (GCP3-GFP, yellow), CAMSAP3 colocalization (magenta) with γ-TuRC-anchored microtubule minus-end and subsequent microtubule release from γ-TuRC in conditions described in **a** with CAMSAP3. Thin arrows indicate γ-TuRC, while thick arrows indicate CAMSAP3. **f,** Still frames at indicated time points in min (left) and kymographs (right) of a 10 min time-lapse video showing microtubule nucleation and rapid microtubule release from γ-TuRC under conditions shown in **e**. **g,h,** Colocalization frequency of γ-TuRC-anchored microtubule minus-ends with CAMSAP3, from experiments shown in **e** (n=5, m=18, l=102 for GCP3-GFP) and **k** (n=5, m=16, l=47 for GCP6-GFP) (**g**). Individual data points represent single experiments from which % of γ-TuRC-anchored microtubule minus-ends colocalizing with CAMSAP3 were quantified, p=0.3157. **h,** Quantification of % of microtubules released from γ-TuRC colocalizing with CAMSAP3, as shown in **g**. n=4, m=18, l=32 for GCP3-GFP and n=5, m=16, l=17 for GCP6-GFP. Individual data points represent single experiments from which % of growing microtubule minus-ends that were labeled with γ-TuRC and CAMSAP3 were quantified, p=0.2065. In **g** and **h**, n is the number of independent experiments, m is the number of fields of view and l is the total number of active γ-TuRCs in **g** and number of active γ-TuRCs colocalizing with CAMSAP3 in **h,** analyzed over 10 min duration. The plots present mean ± s.e.m. Two-tailed unpaired t-tests were used to test for significance. ns, not significant. **i,** Histogram showing frequency distribution of γ-TuRC dissociation times quantified as time interval between the initial CAMSAP3 binding and the onset of microtubule minus-end elongation. n=14 γ-TuRC dissociation events from experiments shown in **e** and pooled from three independent experiments. **j,** Comparison of fluorescence intensities of total active GCP3-GFP molecules nucleating microtubules (n=80); active GCP3-GFP molecules that did not recruit CAMSAP3 (n=52, ns, p=0.9846; active GCP3-GFP molecules that recruited CAMSAP3 and remained attached to the microtubule (n=15, ns, p=0.2382); and active GCP3-GFP molecules that recruited CAMSAP3 and subsequently released microtubule (n=14, ns, p=0.9427). Data was analyzed from experiments shown in **e** and pooled from four independent experiments. One-way ANOVA with Tukey’s multiple comparisons test was used to compare the means with each other. ns, p (magenta)=0.2463. **k,l,** Still frames (at indicated time points in min) (**k**) and representative kymographs (**l**) from a 10 min time-lapse video showing CAMSAP3 colocalization (magenta) with γ-TuRC-anchored microtubule minus-end and subsequent microtubule (cyan) release from γ-TuRC (GCP6-GFP, yellow) in the same conditions as in **a**. Thin arrows indicate γ-TuRC signal, while thick arrows indicate CAMSAP3 signal.

Recent work has shown that also partial/incomplete γ-TuRCs can nucleate microtubules ^22^. To test whether CAMSAP3 preferentially binds to minus ends anchored by incomplete γ-TuRCs and triggers their detachment, we have measured fluorescence intensity of γ-TuRCs in the GFP channel, because it should directly reflect the number of GCP3-GFP molecules. We found the fluorescence intensity of the total active γ-TuRC population was similar to that of γ-TuRCs that nucleated microtubules and did not recruit CAMSAP3 and was not different from those that subsequently recruited CAMSAP3 and either stayed attached or got released (Fig. 2j). To further validate our result, we also generated γ-TuRC that were purified using a homozygous knock-in HEK293T cell line where the GCP6 subunit of γ-TuRC was C-terminally tagged with GFP and Twin-Strep-tag (Extended data Fig. 3g,h). Mass spectrometry showed that γ-TuRC purified from this cell line was similar in terms of components and associated proteins to that purified using GCP3-GFP-SII (Extended data Fig. 3i and Supplementary Table 5). It was shown previously that a subcomplex containing GCP4, 5 and 6 is by itself nucleation-incompetent and needs to be supplemented with cell extracts containing GCP2 and 3 to nucleate microtubules^41^. Therefore, GCP6-containing γ-TuRCs that can nucleate microtubules are likely to be complete rings. In our reconstitutions, we observed that GCP6-GFP-SII-containing γ-TuRCs could nucleate microtubules, which then could recruit CAMSAP3 to their minus ends, initiate minus-end growth and detach from γ-TuRC, and the frequency of CAMSAP3 binding and minus end growth was even slightly higher than for γ-TuRCs purified using GCP3-GFP-SII (Fig. 2g,h,k,l and Supplementary Video 1, right panel). We conclude that CAMSAP3 can bind to the minus ends of a subset of microtubules nucleated and anchored by γ-TuRC, promote minus end polymerization and trigger their release.

### CDK5RAP2 inhibits CAMSAP binding to the minus ends of γ-TuRC-anchored microtubules

To test whether nucleation-promoting factors affect CAMSAP3 binding and microtubule release from γ-TuRC, we first examined whether they were active in the same assay conditions. We found that whereas CDK5RAP2 and CLASP2 could still potentiate γ-TuRC-dependent microtubule nucleation up to ∼20% and ∼9% respectively in the presence of 80 mM KCl and 25 µM tubulin, this was not the case for chTOG (Fig. 3a,b). CAMSAP3 binding to γ-TuRC-anchored minus ends and microtubule release were observed in the presence of either of the three proteins (Fig. 3c-e and Supplementary Video 2). Interestingly, CDK5RAP2, but not CLASP2 or chTOG, strongly suppressed CAMSAP3 binding to the minus ends of γ-TuRC-nucleated microtubules (Fig. 3f). Microtubule minus ends that did recruit CAMSAP3 started to grow and detached from γ-TuRC with a comparable frequency in all conditions (Fig. 3g), though the data for chTOG were less reliable because the combination of high ionic strength and chTOG made microtubule growth events very short-lived and thus limited the time when microtubule release could be observed. Altogether, CDK5RAP2 strongly suppressed CAMSAP3-driven microtubule detachment from γ-TuRC, while this was not the case for CLASP2, and no strong conclusions could be made for chTOG (Fig. 3h).

**Fig. 3.**
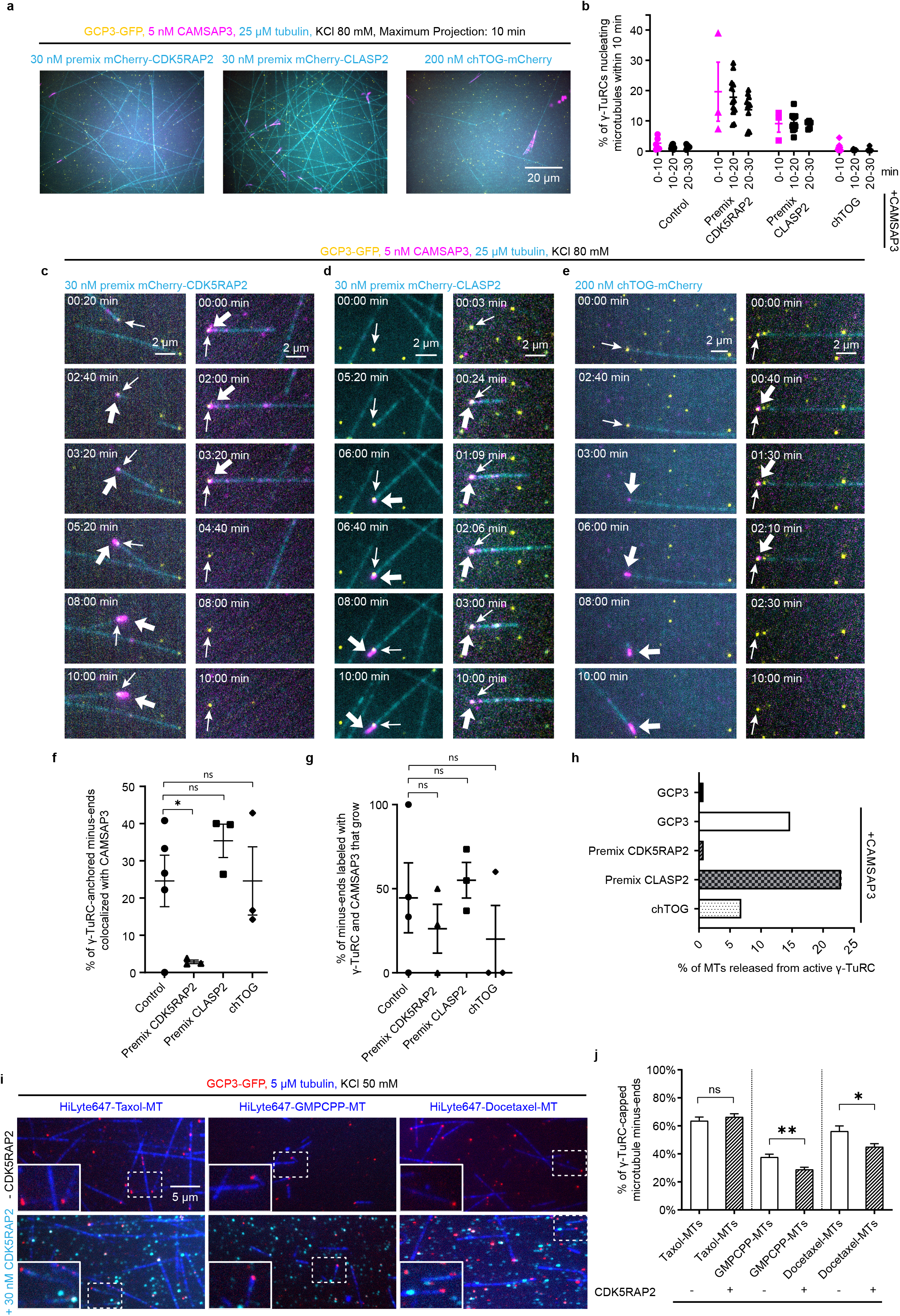
CDK5RAP2 inhibits CAMSAP3 binding to the minus ends of γ-TuRC-anchored microtubules. **a,** Maximum intensity projections of 10 min time-lapse videos, acquired after 10 min of incubation, showing microtubules (cyan) nucleated from γ-TuRC (GCP3-GFP, yellow) in the presence of 25 μM tubulin (24.5 μM unlabeled porcine tubulin and 0.5 μM rhodamine-tubulin), 80 mM KCl and 5 nM SNAP-AF647-CAMSAP3 (magenta) supplemented with either mCherry-CDK5RAP2 (cyan) (preincubated with γ-TuRC); or mCherry-CLASP2 (cyan) (preincubated with γ-TuRC); or 200 nM chTOG-mCherry (cyan). **b,** Plot showing average microtubule nucleation efficiency of γ-TuRC quantified from experiments shown in **a**. For premix CDK5RAP2: n=3 for 0-10 min; n=12 for 10-20 min; n=11 for 20-30 min; analyzed from three independent experiments. For premix CLASP2: n=3 for 0-10 min; n=10 for 10-20 min; n=5 for 20-30 min; analyzed from three independent experiments. For chTOG: n=7 for 0-10 min; n=6 for 10-20 min; n=6 for 20-30 min; analyzed from three independent experiments. n is the number of fields of view analyzed. Control data have been taken from Fig. 2b and replotted here for direct comparison. The plot presents mean ± s.e.m., and each data point represents a single field of view from which % of γ-TuRCs nucleating microtubules were quantified for the given time point from an individual experiment. Data points in magenta (0-10 min) were acquired from a smaller field of view and are not directly comparable to the data points in black (10-20 min and 20-30 min) acquired from a bigger field of view shown in **a**. **c-e,** Still frames (at indicated time points in min) from 10 min time-lapse videos showing two different examples (left and right) of γ-TuRC-CAMSAP3 interplay at the γ-TuRC-anchored microtubule minus-ends under indicated experimental conditions, also shown in **a**. Example on the left illustrates microtubule (cyan) release from γ-TuRC (yellow, thin arrows) colocalizing with CAMSAP3 (magenta, thick arrows) in the presence of indicated proteins (cyan). Example 2 on the right illustrates occasions when CAMSAP3 binds, but does not displace γ-TuRC from the microtubule minus-end. **f,g,** Colocalization frequency of γ-TuRC-anchored microtubule minus-ends with CAMSAP3, from experiments shown in **a,c-e** (**f**). n=3, m=26, l=393 for premix mCherry-CDK5RAP2, *p=0.0394; n=3, m=13, l=310 for premix mCherry-CLASP2, ns, not significant, p=0.2678; and n=3, m=23, l=44 for chTOG-mCherry, ns, p=0.9996. Individual data points represent single experiments from which % of γ-TuRC-anchored microtubule minus-ends colocalizing with CAMSAP3 were quantified. **g,** Quantification of % of microtubules released from the CAMSAP3 colocalized population of γ-TuRC shown in **f**. n=3, m=26, l=12 for premix mCherry-CDK5RAP2, ns, not significant, p=0.4778; n=3, m=13, l=114 for premix mCherry-CLASP2, ns, p=0.6836; and n=3, m=23, l=9 for chTOG-mCherry, ns, p=0.3480. Individual data points represent single experiments from which % of growing microtubule minus-ends that were labeled with γ-TuRC and CAMSAP3 were quantified. In **f** and **g**, n is the number of independent experiments, m is the number of fields of view and l is the total number of active γ-TuRCs in **f** and number of active γ-TuRCs colocalized with CAMSAP3 in **g**, analyzed over 10 min duration. Control data has been taken from Fig. 2g for **f** and from Fig. 2h for **g** (GCP3 values) and replotted here for direct comparison. The plots present mean ± s.e.m. One-way ANOVA with uncorrected Fisher’s LSD tests were used to compare the means with control. **h,** Plot showing frequency of microtubule release from active γ-TuRC in the presence of 5 nM SNAP-AF647-CAMSAP3 under the experimental conditions shown in **a,c-e**. n=393 for premix mCherry-CDK5RAP2, pooled from three independent experiments; n=310 for premix mCherry-CLASP2, pooled from three independent experiments; and n=44 for chTOG-mCherry, pooled from three independent experiments; where n is the number of active γ-TuRCs analyzed. The two control data values (GCP3 with or without CAMSAP3) have been taken from Fig. 2d (GCP3 values with or without CAMSAP3) and replotted here for direct comparison. **i,** Maximum intensity projections of 5 min time-lapse videos showing γ-TuRC-capped minus ends of stabilized microtubules (blue), as indicated, that contain different protofilament number: rhodamine-labeled Taxol-stabilized microtubules with 12 or 13 protofilaments (left) and rhodamine-labeled GMPCPP- and Docetaxel-stabilized microtubules with predominantly 14 protofilaments (middle and right respectively) were incubated with γ-TuRC (20 nM GCP3-GFP, red) in the presence of 5 μM tubulin (4.5 μM unlabeled porcine tubulin and 0.5 μM rhodamine-tubulin) and 50 mM KCl with (bottom) or without (top) 30 nM SNAP-AF647-CDK5RAP2 (cyan). Insets show enlarged view of the ROI marked with dashed white line. **j,** Plot generated from experiments shown in **i**, comparing minus-end capping efficiency of γ-TuRC for stabilized microtubules with different protofilament numbers in the absence or presence of 30 nM SNAP-AF647-CDK5RAP2. n=18, m=1075, N=3 for Taxol-stabilized microtubules without CDK5RAP2; n=11, m=1420, N=3 for Taxol-stabilized microtubules with CDK5RAP2; n=29, m=3032, N=3 for GMPCPP-stabilized microtubules without CDK5RAP2; n=26, m=2454, N=3 for GMPCPP-stabilized microtubules with CDK5RAP2; n=11, m=1253, N=3 for Docetaxel-stabilized microtubules without CDK5RAP2; n=15, m=1171, N=3 for Docetaxel-stabilized microtubules with CDK5RAP2; where n is the number of fields of view analyzed, m is the number of microtubule minus-ends analyzed and N is the number of independent experiments. The bar plot presents mean ± s.e.m., and individual data points represent single fields of view from which % of minus-ends capped by γ-TuRC were quantified. One-way ANOVA with Šídák’s multiple comparisons test was used to compare the pre-selected pair of means. ns, not significant, p=0.8573; *p=0.0187; **p=0.0050.

We next hypothesized that CDK5RAP2 exerts its effect by altering γ-TuRC geometry so that it would be more similar to a 13-protofilament microtubule, which would be beneficial for microtubule nucleation and at the same time inhibit CAMSAP3 binding. We first tested this possibility by performing negative stain transmission EM of γ-TuRC, either alone or incubated in the presence of 120 nM CDK5RAP2 but found no support for this idea (Extended data Fig.4). γ-TuRC alone in negatively stained micrographs and 3D reconstructions of density maps looked similar to previously reported work and fitted well into the density map of a published model (from PDB ID: 6V6S ^20^) (Extended data Fig. 4a-c,i). However, we could not detect any noticeable differences in the γ-TuRC structure in the presence (Extended data Fig. 4d-f,j) and absence of CDK5RAP2 (Extended data Fig. 4g). Both reconstructions deviated significantly from the closed conformation of γ-tubulin small complex (γ-TuSC) oligomers (from PDB ID: 5FLZ ^42^), and did not match the microtubule geometry (from PDB ID: 2HXF ^43^ and EMD-5193 ^44^) (Extended data Fig. 4h). It should be noted that the densities of terminal γ-TuSC at the γ-TuRC seam and in the luminal bridge were not clearly resolved in our γ-TuRC reconstructions. Therefore, we cannot make conclusions about potential conformational changes that could occur at these sites.

As an alternative approach, we used a microtubule-capping assay, whereby γ-TuRCs bind to the ends of preformed microtubules. We have compared Taxol-stabilized microtubules, which in our hands have 12 or 13 protofilaments to GMPCPP- and Docetaxel-stabilized microtubule preparations, which predominantly have 14 protofilament microtubules ^45^. As expected, Taxol-stabilized microtubules were capped by γ-TuRC more efficiently than Docetaxel-stabilized ones, and GMPCPP-stabilized were capped even less well, possibly because they have more curved terminal protofilaments than the taxane-stabilized ones ^46, 47^ (Fig. 3i,j). Interestingly, the addition of CDK5RAP2 significantly inhibited capping of both types of 14-protofilament microtubules, without affecting the capping efficiency of Taxol-bound microtubules (Fig. 3i, j). These data are in line with the idea that in the presence of microtubules, CDK5RAP2 promotes a conformational change or conformational flexibility in γ-TuRC that would allow a better match with the 13-protofilament microtubule geometry.

### CAMSAPs cause γ-TuRC detachment by stabilizing growing microtubule minus ends

Since in our assays microtubules rarely dissociated from γ-TuRC spontaneously, their CAMSAP3-induced detachment must be an active process. To get more insight into this process, we compared the effects of CAMSAP2 and CAMSAP3, which decorate and stabilize growing microtubule minus ends, and CAMSAP1, which labels free minus ends but does not decorate or stabilize them^29^. All three CAMSAPs could bind to γ-TuRC-anchored minus ends, with CAMSAP2 being slightly less efficient (Fig. 4a-c and Extended data Fig. 3d). However, although CAMSAP1 and CAMSAP3 bound to the minus ends equally well, CAMSAP1 had little effect on microtubule release from γ-TuRC (Fig. 4d,e and Supplementary Video 3). These data support the view that γ-TuRC is displaced from the minus ends due to their elongation, which is made more robust by CAMSAP2 and CAMSAP3 that stably decorate the newly formed microtubule lattice and may also alter its conformation.

**Fig. 4.**
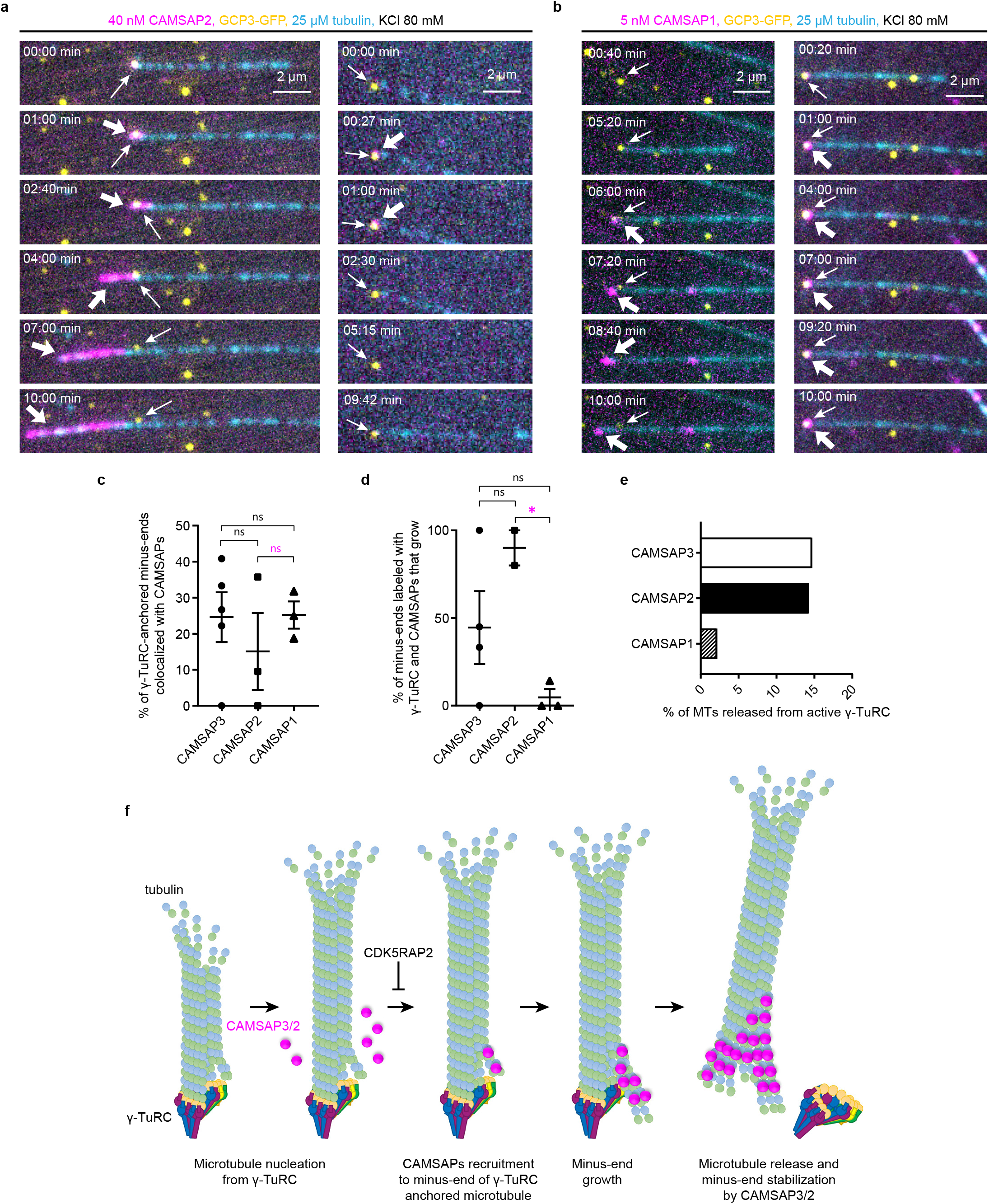
CAMSAPs cause γ-TuRC detachment by stabilizing growing microtubule minus ends. **a,b,** Still frames (at indicated time points in min) from 10 min time-lapse video showing two different examples (left and right) of γ-TuRC interplay with CAMSAP2 (**a**) or CAMSAP1 (**b**) in the indicated conditions in the presence of 25 μM tubulin (24.5 μM unlabeled porcine tubulin and 0.5 μM rhodamine-tubulin). Example on left illustrates microtubule release; example on right illustrates occasions when CAMSAP2 (**a**) or CAMSAP1 (**b**) does not displace γ-TuRC from microtubule minus-ends. **c,d,** Colocalization frequency of γ-TuRC-anchored microtubule minus-ends with CAMSAPs, from experiments shown in **a,b** (**c**). n=3, m=15, l=42 for CAMSAP2, ns, not significant, p=0.4000; and n=3, m=18, l=46 for CAMSAP1, ns, p=0.9581. ns, p (magenta)=0.4234. Individual data points represent single experiments from which % of γ-TuRC-anchored microtubule minus-ends colocalized with CAMSAPs were quantified. **d,** Quantification of % of microtubules released from γ-TuRCs colocalizing with CAMSAPs, as shown in **a,b**. n=3, m=15, l=7 for CAMSAP2, ns, p=0.1346; and n=3, m=18, l=12 for CAMSAP1, ns, p=0.1365. *p (magenta)=0.0217. Individual data points represent single experiments from which % of growing microtubule minus-ends that were labeled with γ-TuRC and CAMSAPs were quantified. In **c** and **d**, n is the number of independent experiments, m is the number of fields of view and l is the total number of active γ-TuRCs in **c** and number of active γ-TuRCs colocalized with CAMSAPs in **d**, analyzed over 10 min duration. Control data (CAMSAP3 values) have been taken from Fig. 2g for **c** and from Fig. 2h for **d** (GCP3 values) and replotted here for direct comparison. The plots present mean ± s.e.m. One-way ANOVA with uncorrected Fisher’s LSD tests were used to compare the means with each other. **e,** Plot showing frequency of microtubule release from active γ-TuRC in the presence of CAMSAPs under the experimental conditions shown in **a,b**. n=42 for CAMSAP2, pooled from three independent experiments; n=46 for CAMSAP1, pooled from three independent experiments; where n is the number of active γ-TuRCs analyzed. Control data (CAMSAP3) has been taken from Fig. 2d (GCP3 values with CAMSAP3) and replotted here for direct comparison. **f,** Model: A microtubule is first nucleated by γ-TuRC with some protofilaments that are not attached to the γ-tubulin subunits, allowing them to attain a conformation permissive for CAMSAP binding. CAMSAP2 or 3 binds to the minus-end of the γ-TuRC-capped microtubule at an intradimer site between two protofilaments ^14^, promotes minus-end polymerization and stabilizes the growing minus-end lattice. Growing minus-end pushes away γ-TuRC, and the microtubule is released. CDK5RAP2 inhibits microtubule release, possibly by altering γ-TuRC conformation to fit the geometry of a 13-protofilament microtubule and therefore, suppresses CAMSAP binding.

## Discussion

In this study, we have uncovered a new mechanism of generation of stable microtubule minus ends – displacement of γ-TuRC from a newly nucleated microtubule by a polymerizing minus end, which is promoted by CAMSAP2 and CAMSAP3. γ-TuRC is the major microtubule-nucleating factor in most cell types, but particularly in cells with dense microtubule arrays, many microtubule minus ends depend for their stabilization and anchoring on CAMSAP family members. How CAMSAP-stabilized microtubules are generated is currently unclear. CAMSAPs are diffusely distributed in mammalian cells where microtubules are depolymerized^29^, but can bind to microtubule minus ends released from nucleation centers such as the centrosome or the Golgi^29, 48^. While such release can in principle be induced by severing enzymes or motor proteins pulling on microtubules^49^, here we show that CAMSAPs themselves can be sufficient to mediate this process. The microtubule minus-end binding domain of CAMSAPs, CKK, recognizes the minus ends by binding to an interface between two curved protofilaments^14^. Such protofilament arrangement would not occur at microtubule minus ends fully attached to a γ-TuRC^16, 17^. However, given the asymmetric γ-TuRC structure^18–21^ and the fact that not all γ-tubulin subunits within γ-TuRC are essential for microtubule nucleation^22^, it is possible that some protofilaments at the γ-TuRC-capped minus end will be unattached and acquire a flared conformation permissive for CAMSAP binding (Fig. 4f). Having bound to some sites at the interface between the microtubule minus end and γ-TuRC, CAMSAPs could stabilize free protofilaments and promote their elongation. Furthermore, CAMSAPs could potentially trigger detachment of neighboring protofilaments from γ-TuRC by inducing protofilament skew^14^. Protofilaments elongating from the minus end would be expected to generate a pushing force, similar to growing microtubule plus ends ^50^, that would contribute to microtubule release from γ-TuRC. In accordance with this model, stable decoration of minus-end-grown lattice, a property common for CAMSAP2 and CAMSAP3 but not CAMSAP1^29^ is required for release. Repeated activity of γ-TuRC in the presence of CAMSAP2 or CAMSAP3 thus can lead to generation of a pool of stable microtubule minus ends that are not directly attached to their nucleation sites. Such a “handover” mechanism can help to increase microtubule density in different settings. For example, it can generate free, stable non-centrosomal minus ends in cells such as fibroblasts or cancer cells, or amplify microtubules at non-centrosomal MTOCs such as the Golgi apparatus or the apical cortex of epithelial cells, where γ-TuRC and CAMSAPs operate in parallel^1^. Moreover, a pool of CAMSAP-stabilized minus ends could recruit katanin for severing and further amplification of non-centrosomal microtubules^29, 49^.

Similar to microtubule nucleation, microtubule release from γ-TuRC is expected to be tightly regulated in order to control the abundance and positioning of microtubule minus ends. Our data suggest that the factors promoting microtubule nucleation can play a role in this process. In this study, we focused on three such factors, chTOG, CLASP2 and CDK5RAP2. While we were able to confirm that chTOG potently promotes γ-TuRC-dependent microtubule nucleation, the incompatibility in experimental conditions did not allow us to reach firm conclusions about its interplay with CAMSAPs. However, we were successful in examining the effect of CDK5RAP2. This protein is well-known for its ability to bind γ-TuRC through the domain called γ-TuNA (γ-TuRC nucleation activator), but there was some conflicting evidence about its ability to activate γ-TuRC in vitro^19, 26, 27, 34, 51^. Two studies showed a stimulatory effect of nanomolar γ-TuNA concentrations on microtubule nucleation by γ-TuRC purified from mammalian cells using its affinity for γ-TuNA ^26, 27^, whereas, more recently, two other groups found no to little effect of even micromolar γ-TuNA concentrations on *Xenopus* γ-TuRC when purified using a γ-tubulin antibody^19, 51^. A recent study attributed these differences to the presence of bulkier N-terminal tags on γ-TuNA and purification methods involving different affinity tags^34^.

Here, we show that full-length CDK5RAP2 can stimulate γ-TuRC-dependent microtubule nucleation already at 30 nM concentration, which is consistent with previously published data^26^. Interestingly, CDK5RAP2 also suppressed CAMSAP binding to γ-TuRC-anchored minus ends and as well as capping of 14- but not 13-protofilament microtubules. All these data can be best explained by the ability of CDK5RAP2 to trigger a conformational change that would make γ-TuRC match better with the 13-protofilament microtubule geometry. However, we could find no structural support for this idea. Since the densities of terminal γ-TuSC (GCP2_13_ and GCP3_14_) and the luminal bridge were not clearly resolved in our reconstructions, so we cannot exclude the possibility of CDK5RAP2-induced conformational changes occurring at these sites. Notably, the terminal γ-TuSC constitutes the binding site for a dimeric γ-TuNA^52^. In a recent comparative study, γ-TuNA-bound human γ-TuRC showed a structural difference at the interface of GCP6 and neighboring GCP2 when compared to that of γ-TuNA-unbound *Xenopus* γ-TuRC^23^. Furthermore, it is also possible that CDK5RAP2 might only contribute to a local conformational change in the terminal γ-TuSC^23^, and this small change would synergize with other γ-TuRC activation mechanisms such as binding of tubulin dimers or activators^6, 7, 51^. Further study would be needed to find out whether the lack of detectable CDK5RAP2-induced structural change in our reconstructions was due to technical reasons or because this alteration is manifested as increased flexibility, is transient and/or requires binding to a microtubule.

We also discovered that γ-TuRC-dependent microtubule nucleation can be strongly enhanced by CLASP2, a positive regulator of microtubule density and growth^53^. Given the abundant colocalization between CLASP2 and γ-TuRC even in the absence of free tubulin, they might bind to each other directly. Previous work has shown that CLASP2 potently promotes formation of complete tubes from incomplete tubulin assemblies and microtubule outgrowth from seeds at low tubulin concentration^28, 54^. The mechanism underlying these activities needs further study, but since CLASP2 did not inhibit CAMSAP binding to γ-TuRC-anchored minus ends and their release, it is unlikely to affect γ-TuRC geometry but more likely acts by affecting microtubule polymerization intermediates. Altogether, our work shows that mechanisms controlling microtubule nucleation and the players involved in this process can also affect the destiny of the generated microtubule minus-ends.

## Author Contributions

D.R. designed and performed protein purifications and in vitro reconstitution experiments, analyzed data and wrote the paper; S.H., J.L.M. and Y.Z. have generated and characterized essential reagents; R.S., K.E.S. and A.F. M.A. performed, analyzed and supervised mass spectrometry experiments; E.A.K. facilitated and performed data analysis; M.W. generated and analyzed transmission EM data; K.J. generated reagents, performed experiments, supervised the project and wrote the paper; A.A. coordinated the project and wrote the paper.

## Acknowledgements

Negative stain EM grid screening was performed with support of the ScopeM imaging center, ETH Zurich. Negative stain EM data collection was performed with support of the Center for Microscopy and Image Analysis, University of Zurich. The authors gratefully acknowledge Lina Carlini for her help in developing the MATLAB analysis scripts used to compare γ-TuRC structural data. This work was supported by the European Research Council Synergy grant 609822 and the ZonMW TOP 91216006 program to A.A and grants from National Natural Science Foundation of China (31871356, 32070705) to K.J.

## Methods

### DNA constructs

We used previously described SII-mCherry-CLASP2 construct^28^. chTOG construct was a gift of S. Royle (University of Warwick, UK). chTOG-mCherry-SII was made by cloning the full length construct in a modified pTT5 expression vector (Addgene no. 44006) with a C-terminus mCherry-Twin-Strep-tag. GFP-CDK5RAP2 was a gift from Robert Z. Qi (The Hong Kong University of Science and Technology, China). SII-mCherry-CDK5RAP2, SII-SNAP-CAMSAP3, Bio-Tev-mCherry-CAMSAP2 and SII-SNAP-CAMSAP1 were made by cloning the full length previously described constructs^29^ in modified C1 vectors with either a Twin-Strep-mCherry, or Twin-Strep-SNAP, or Bio-Tev-mCherry tag at the N-terminus.

### Cell lines and cell culture

HEK293T cells (from ATCC) were cultured in Dulbecco’s Modified Eagle’s Medium DMEM/Ham’s F10 media (1:1) supplemented with 10% fetal calf serum (FCS) and 1% antibiotics (penicillin and streptomycin). The cell line used here was not found in the commonly misidentified cell lines database maintained by the Immunization Coalition of Los Angeles County. No further cell line authentication was performed. The cell lines were routinely checked for mycoplasma contamination using LT07-518 Mycoalert assay. Polyethylenimine (PEI, Polysciences) was used to transfect HEK293T cells with plasmids for StrepTactin- and Streptavidin-based protein purification at 3:1 ratio of PEI:plasmid.

### Generation of homozygous HEK93T knock-in cell lines endogenously tagged with a GFP-SII for GCP3 and GCP6

GCP3-GFP-SII and GCP6-GFP-SII knock-in cell lines were generated using CRISPR-Cas9 technology^55^. To generate knock-in cell lines, CRISPR guide RNAs were designed using the web tool from MIT: http://crispr.mit.edu/. The guide RNAs were designed to overlap with the stop-codon to disrupt the recognition site after insertion of the tag avoiding any further cleavage. Annealed oligo’s were inserted into pSpCas9(BB)-2A-Puro (px459, Addgene #62988) using BbsI. Donor plasmids were designed by selecting 800-1000 bp of homology flanking both sides of the stop codon of the targeted gene. The two homology arms were obtained by genomic DNA PCR from HeLa cells. GFP-SII tag was amplified by PCR using primers with complementary domains for the homology arms. Using Gibson assembly, the two homology arms and GFP-SII were cloned into the donor plasmid. FuGENE6 (Roche) was used to cotransfect cells with px459 containing humanized Cas9, guide RNA followed by tracrRNA and a puromycin resistance marker together with a donor construct. 24 hours post-transfection, cells were selected for two days using 2 μg/mL Puromycin and subsequently subcloned to a single cell dilution. Positive clones were confirmed using immunofluorescence, genomic DNA-PCR genotyping and Western blotting. Following are the guide RNA sequences and primers used: for GCP3, gRNA, 5’-GGACCGCGAGCTTCACGTGT-3’; 5’-homology arm, 5’- TCAACACAGCAGAGCCTGTGC-3’ and 5’-CGTGTGGGAGCTGCGCCGCC-3’; 3’-homology arm, 5’-AGCTCGCGGTCCTCCCAGGG-3’ and 5’-CGAATGCATCTGAAAGATAATTGC-3’; genotyping, 5’-GGAAGGAAAAACAGACCCAACC-3’ and 5’- CGAATGCATCTGAAAGATAATTGC-3’. For GCP6, gRNA, 5’- CAGAGCAGCCTCAGGCGTCC-3’; 5’-homology arm, 5’-TTTCTGCCTAGCTTGGAGCTG-3’ and 5’-GGCGTCCTGGTAGTAGTTGTTGAAGTTG-3’; 3’-homology arm, 5’- GGCTGCTCTGCGGGGGAC-3’ and 5’-CTACAGGCGTACAGGTGAGC-3’.

### Immunofluorescence and Western blotting

HEK293T GCP3- and GCP6-GFP-SII knock-in cells, seeded on coverslips, were fixed with prechilled methanol at -20°C for 10 min followed by three washes with PBS and mounting on glass slides in Vectashield mounting medium containing DAPI (Vector laboratories).

For Western blotting of GCP3- and GCP6-GFP-SII HEK293T cell lysates, cells grown in 6-well plates were harvested and lysed in lysis buffer containing 20 mM Tris-Cl pH 7.5, 100 mM NaCl, 1% Triton-X-100 supplemented with 10% glycerol and complete protease inhibitor cocktail (Roche). Lysates were cleared by centrifugation at 21,000xg for 20 min at 4 ⁰C. 20 μg supernatants from the above step or 35 μg purified γ-TuRC were loaded on 8% SDS-PAGE gels, then transferred onto a nitrocellulose membrane (Sigma-Aldrich). Membranes were blocked in 2% BSA in 0.02% Tween-20 in PBS for 30 min at room temperature followed by overnight incubation with primary antibodies (rabbit polyclonal anti-GFP (1:4000, Abcam, ab290); mouse anti-GCP3 (1:1000, Santa Cruz, sc-373758); mouse monoclonal anti-GCP6 (1:500, Santa Cruz, sc-374063); mouse monoclonal anti-GCP5 (1:500, Santa Cruz, sc-365837); mouse monoclonal anti-GCP2 (1:500, Santa Cruz, sc-377117); rabbit polyclonal anti-GCP4 (1:1000, ThermoFisher, PA5-30557) and mouse monoclonal anti-γ-tubulin (1:10000, Sigma, T6557)) at 4°C, three washes with 0.02% Tween-20 in PBS, 1 hour incubation with secondary antibodies (1:15000, goat anti-rabbit IRDye-800CW and goat anti-mouse IRDye-680LT from Li-Cor Biosciences, Lincoln, LE) at room temperature and final three washes. Membranes were imaged using Odyssey CLx infrared imaging system (Li-Cor Biosciences).

### Purification of γ-TuRC from HEK293T GCP3- and GCP6-GFP-SII knock-in cells

Human γ-TuRCs used in the in vitro reconstitution assays were purified using Twin-Strep-tag and Strep-Tactin affinity purification method as described previously^56^. Homozygous HEK293T GCP3- and GCP6-GFP-SII knock-in cells, cultured in dark, were harvested from eight 15 cm dishes each and resuspended and lysed in lysis buffer (50 mM HEPES, 150 mM NaCl, 0.5% Triton-X-100, 1 mM MgCl_2_, 1 mM EGTA, 0.1mM GTP and 1 mM DTT, pH 7.4) supplemented with EDTA-free protease inhibitor cocktail (Roche). Cell lysates were subjected to centrifugation at 21,000xg for 20 min at 4°C. The supernatants obtained from the previous step were incubated with equilibriated Strep-Tactin Sepharose beads (28-9355-99, GE Healthcare) for 45 min at 4 °C. Following incubation, beads were washed three times with the wash buffer (50 mM HEPES, 150 mM NaCl and 0.1% Triton-X-100, 1 mM MgCl_2_, 1 mM EGTA, 0.1mM GTP and 1 mM DTT, pH 7.4) and γ-TuRC was eluted for 15 min at 4 °C in elution buffer (50 mM HEPES, 150 mM NaCl (300 mM NaCl for GCP6-GFP-SII), 0.05% Triton-X-100, 1 mM MgCl_2_, 1 mM EGTA, 0.1mM GTP, 1 mM DTT and 2.5 mM d-Desthiobiotin, pH 7.4). GCP6-tagged purified γ-TuRC was then subjected to buffer exchange using Vivaspin 500 centrifugal concentrator (10 kDa MWCO, Sartorius VS0102) for a final NaCl concentration of 150 mM in eluate. Purified γ-TuRCs were immediately aliquoted, snap-frozen in liquid N2 and stored at -80 °C. Throughout the purification process, tubes were covered with aluminium foil wherever possible. The purity and composition of purified γ-TuRCs were analyzed by western blot and mass spectrometry.

### Purification of recombinant proteins from HEK293T cells for in vitro reconstitution assays

Human mCherry-CDK5RAP2, mCherry-CLASP2, chTOG-mCherry and SNAP-AF647-CAMSAP1 and mouse SNAP-AF647-CAMSAP3 used in the in vitro reconstitution assays were purified using same Twin-Strep-tag and Strep-Tactin affinity purification method as described above for γ-TuRC, but with modified buffers and steps. In brief, HEK293T cells transfected with 50 μg of respective constructs per 15 cm dish were harvested 36 hours post-transfection from four 15 cm dishes each and resuspended and lysed in lysis buffer (50 mM HEPES, 300 mM NaCl, 0.5% Triton-X-100, 1 mM MgCl_2_ and 1 mM EGTA, pH 7.4) supplemented with EDTA-free protease inhibitor cocktail (Roche). Cell lysates were clarified and the supernatants obtained were incubated with equilibriated Strep-Tactin Sepharose beads. Following incubation of mCherry-CDK5RAP2, mCherry-CLASP2 and chTOG-mCherry preparations, beads were additionally washed five times using high salt (1 M NaCl) containing wash buffer (50 mM HEPES, 0.1% Triton-X-100, 1 mM MgCl_2_ and 1 mM EGTA, pH 7.4) before washing three times with 300 mM NaCl containing wash buffer. For SNAP-tag labeling of CAMSAP3 and CAMSAP1 with Alexa Fluor 647 dye, washed beads were incubated with labeling mix (50 μM Alexa Fluor 647 dye in 50 mM HEPES, 150 mM NaCl and 0.1% Triton-X-100, 1 mM MgCl_2_, 1 mM EGTA and 1 mM DTT, pH 7.4) for 1 hour. Following this incubation, beads were washed five times with wash buffer containing 300 mM NaCl to remove excess dye. Proteins were then eluted in elution buffer containing 50 mM HEPES, 150 mM NaCl, 0.05% Triton-X-100, 1 mM MgCl_2_, 1 mM EGTA, 1 mM DTT and 2.5 mM d-Desthiobiotin, pH 7.4.

For purification of human mCherry-CAMSAP2, HEK293T cells were transfected with 25 μg of Bio-Tev-mCherry-CAMSAP2 and 25 μg of BirA per 15 cm dish. Cells were harvested 36 hours post-transfection from four 15 cm dishes and resuspended in lysis buffer (50 mM HEPES, 300 mM NaCl, 0.5% Triton-X-100, 1 mM MgCl_2_, 1 mM EGTA and 1 mM DTT, pH 7.4) supplemented with EDTA-free protease inhibitor cocktail (Roche). Cell lysate was incubated with Dynabeads M-280 streptavidin (Invitrogen-11206D) for 1 hour. Beads were washed thrice with lysis buffer without protease inhibitors and thrice with the TEV cleavage buffer (50 mM HEPES, 150 mM NaCl, 0.05% Triton-X-100, 1 mM MgCl_2_, 1 mM EGTA and 1 mM DTT). mCherry-CAMSAP2 was eluted in 50 μl TEV cleavage buffer containing 0.5 μg of glutathione S-transferase-6x-histidine Tobacco etch virus protease site (Sigma-Aldrich) for 2 hr at 4 °C.

Purified proteins were immediately aliquoted, snap-frozen in liquid N2 and stored at -80 °C. Bacterially expressed mCherry-EB3 was a gift of Dr. M.O. Steinmetz (Paul Scherrer Institut, Switzerland); they were produced as described previously ^57^. Purity of the samples was analyzed by Coomassie-staining of SDS-PAGE gels and mass spectrometry.

### Mass spectrometry

Purified γ-TuRC preparations (GCP3- and GCP6-tagged) were resuspended in 20 μL of Laemmli Sample buffer (Biorad) and were loaded on a 4%–12% gradient Criterion XT Bis-Tris precast gel (Biorad). Gel was fixed with 40% methanol/10% acetic acid and then stained with colloidal Coomassie dye G-250 (Gel Code Blue Stain Reagent, Thermo Scientific) for 1 hour. Samples were resuspended in 10% formic acid (FA)/5% DMSO post in-gel digestion, and analyzed with an Agilent 1290 Infinity (Agilent Technologies, CA) LC, operating in reverse-phase (C18) mode, coupled to an Orbitrap Q-Exactive mass spectrometer (Thermo Fisher Scientific, Bremen, Germany). Onto a trap column (Reprosil C18, 3 μm, 2 cm x 100 μm; Dr. Maisch) with solvent A (0.1% formic acid in water), peptides were loaded at a maximum pressure of 800 bar and chromatographically separated over the analytical column (Zorbax SB-C18, 1.8 μm, 40 cm x 50 μm; Agilent) using 90 min linear gradient from 7%–30% solvent B (0.1% formic acid in acetonitrile) at a flow rate of 150 nL/min. The mass spectrometer automatically switched between MS and MS/MS in a data-dependent acquisition (DDA) mode. The 10 most abundant peptides were subjected to HCD fragmentation after a survey scan from 350-1500 m/z. MS spectra in high-resolution mode (R > 30,000) were acquired, whereas MS2 was in high-sensitivity mode (R > 15,000). Raw data were processed using Proteome Discoverer 1.4 (version 1.4.0.288, Thermo Scientific, Bremen, Germany) and a database search was performed using Mascot (version 2.4.1, Matrix Science, UK) against a Swiss-Prot database (taxonomy human). Whereas oxidation of methionine was set as a variable modification, carbamidomethylation of cysteines was set as a fixed modification. Up to two missed cleavages were allowed by Trypsin. Data filtering performed using percolator resulted in 1% false discovery rate (FDR). Additional filters set were search engine rank 1 and mascot ion score >20.

To assess the quality of purified recombinant proteins (mCherry-CDK5RAP2, mCherry-CLASP2 and chTOG-mCherry), samples were digested using S-TRAP micro filters (ProtiFi) according to the manufacturer’s protocol. In brief, 4 μg of protein samples were denatured using 5% SDS buffer and reduced and alkylated using DTT (20 mM, 10 min, 95 °C) and iodoacetamide (IAA, 40 mM, 30 min), followed by acidification and precipitation using a methanol triethylammonium bicarbonate (TEAB) buffer before finally loading on a S-TRAP column. Trapped proteins were washed four times with methanol TEAB buffer followed by overnight Trypsin (1 μg, Promega) digestion at 37 °C. Before LC-MS analysis, digested peptides were eluted and dried in a vacuum centrifuge.

Samples were analyzed by reversed phase nLC-MS/MS using an Ultimate 3000 UHPLC coupled to an Orbitrap Q Exactive HF-X mass spectrometer (Thermo Scientific). Digested peptides were separated over a 50 cm reversed phase column, packed in-house, (Agilent Poroshell EC-C18, 2.7 μm, 50 cm x 75 μm) using a linear gradient with buffer A (0.1% FA) and buffer B (80% acetonitrile, 0.1% FA) ranging from 13-44% B over 38 min at a flow rate of 300 nL/min, followed by a column wash and re-equilibration step. MS data was acquired using a DDA method with the set MS1 scan parameters in profile mode: 60,000 resolution, automatic gain control (AGC) target equal to 3E6, scan range of 375-1600 m/z, maximum injection time of 20 ms. The MS2 scan parameters were set at 15,000 resolution, with an AGC target set to standard, an automatic maximum injection time and an isolation window of 1.4 m/z. Scans were acquired using a fixed first mass of 120 m/z and a mass range of 200-2000 and a normalized collision energy (NCE) of 28. Precursor ions were selected for fragmentation using a 1 second scan cycle, a 10 s dynamic exclusion time and a precursor charge selection filter for ion possessing +2 to +6 charges. The total data acquisition time was 55 min.

Raw files were processed using Proteome Discoverer (PD) (version 2.4, Thermo Scientific). A database search was performed for MS/MS fragment spectra using Sequest HT against a human database (UniProt, year 2020) that was modified to include protein sequences from our cloned constructs and a common contaminants database. A precursor mass tolerance of 20 ppm and a fragment mass tolerance of 0.06 Da was set for the search parameters. Up to two missed cleavages were allowed by Trypsin digestion. Carbamidomethylation was set as fixed modification and methionine oxidation and protein N-term acetylation was set as variable modifications. Data filtering performed using percolator resulted in 1% FDR for peptide spectrum match (PSM) and a 1% FDR was applied to peptide and protein assemblies. An additional filter with a minimum Sequest score of 2.0 was set for PSM inclusion. MS1 based quantification was performed using the Pecursor Ion Quantifier node with default settings and precursor ion feature matching was enabled using the Feature Mapper node. Common protein contaminants were filtered out from the results table.

### Negative-stain EM and data processing

Purified γ-TuRC was thawed and incubated in the presence or absence of 120 nM CDK5RAP2 on ice for 20 min. 2-3 μl of protein was applied to glow-discharged carbon-coated copper grids (EMS; CF-400-Cu) and incubated for 45 s at room temperature. Protein solution was removed by manual blotting with Whatman No. 1 filter paper. Then 2-3 μl of protein solution was applied again to improve particle density. The protein solution was manually blotted from one side of the grid while freshly filtered 1% uranyl acetate (wt/vol) was simultaneously pipetted from the opposite side to exchange the solution. Grids were incubated in uranyl acetate for a further 45 s. Stain was removed by manual blotting, and grids were air-dried for >24–48 h in a sealed container containing desiccant before imaging. Grids were initially screened on a TFS Tecnai F20 located at ETH Zurich’s ScopeM facility, and final datasets on suitable grids were collected on an FEI Talos 120 at the University of Zurich’s Center for Microscopy and Image Analysis.

For each condition, several thousand micrographs were recorded via the MAPS automated acquisition software at a magnification of 57,000 X (2.4 Å/pixel) on a BM-Ceta CMOS camera. Contrast transfer function (CTF) parameters were estimated using CTFFIND4 ^58^. All subsequent processing was done in RELION version 3.1 and UCSF Chimera and ChimeraX ^59–61^.

The main 3D reconstruction workflow steps are outlined in Extended data Fig. 4i,j. Generally, micrographs were imported, and a small set of <1,000 particles was manually picked and subjected to reference-free 2D classification. The resulting set of averages was used as an initial template for RELION’s built-in auto-picking implementation. One or two rounds of auto-picking were performed to yield the best templates for optimal picking. Auto-picked particles were binned by four and subjected to reference-free 2D classification to remove particles likely corresponding to dirt and other contaminants. A random subset of the cleaned, binned particles was used to generate an ab initio model. Then, all cleaned, binned particles were subjected to an intermediate 3D auto-refinement step using the ab initio model as a reference, re-extracted using the refined coordinates, and subjected to 3D classification. Particles that generated 3D classes containing the highest level of detail were re-extracted from micrographs at either bin-2 pixel size (4.8 Å) and subjected to a final round of 3D auto-refinement using one of the classes as a new reference model.

A published model for γ-tubulin bound to GCP2 (PDB ID: 6V6S ^20^) was individually docked into each radial “spoke” density of the resulting EM density maps using UCSF Chimera’s “Fit in map” function. Only resulting fits for which i) the number of atoms outside the contour was <20% of the total number of atoms fitted and ii) the resulting fit did not form domain clashes with neighbouring γ-tubulin/GCP2 subunits were kept for further analysis. This procedure led to 12 out of possible 14 γ-tubulin/GCP2 subcomplexes to be reliably fitted into both density maps (Extended data Fig. 4b,c,e,f).

A model for twelve laterally-associated *β*-tubulin subunits was constructed using PDB ID: 2HXF ^43^ and EMD-5193^44^. The ring was aligned to each γ-tubulin ring using the align command in Pymol; the 6th γ-tubulin subunit was used as an alignment anchor. The center of mass of each *β*- or γ-tubulin subunit was then calculated based on its C*α* coordinates and the radial or axial displacement relative to the helical axis of the *β*-tubulin ring (i.e., the microtubule) was determined using a custom MATLAB script. A similar analysis was performed using the γ-tubulin ring from the γ-TuSC oligomer in the “closed” conformation (blue; PDB ID: 5FLZ ^42^). The results are plotted in Extended data Fig. 4h; values close to zero indicate a closer fit to the 13-protofilament microtubule.

### In vitro reconstitution assays

#### Stabilized microtubule preparations

Taxol- and Docetaxel-stabilized and double-cycled GMPCPP-stabilized microtubules used for in vitro γ-TuRC capping assays were prepared as described previously ^45, 62^. In brief, GMPCPP-stabilized microtubules seeds were prepared in the presence of GMPCPP (Guanylyl-(a,b)-methylene-diphosphonate (Jena Biosciences)) by two rounds of polymerization and a depolymerization cycle. First, a 20 μM porcine brain tubulin (Cytoskeleton) mix composed of 70% porcine unlabeled tubulin, 18% biotin tubulin and 12% HiLyte647-tubulin was incubated with 1 mM GMPCPP in MRB80 buffer (pH 6.8, 80 mM K-PIPES, 1 mM EGTA and 4 mM MgCl_2_) at 37°C for 30 minutes. Ploymerization mix was then pelleted by centrifugation in an Airfuge for 5 min at 119,000xg followed by resuspension and depolymerization in MRB80 buffer on ice for 20 min and subsequent polymerization for 30 min in presence of fresh 1 mM GMPCPP at 37°C. GMPCPP-stabilized microtubule seeds were pelleted, resuspended in MRB80 buffer containing 10% glycerol, aliquoted, snap-frozen in liquid N2 and stored at -80 °C.

Taxol- and Docetaxel-stabilized microtubules were prepared 24 hours in advance by polymerizing a 29 μM porcine brain tubulin mix (86% porcine unlabeled tubulin, 10% biotin tubulin and 4% HiLyte647-tubulin) in the presence of 2.5 mM GTP (Sigma-Aldrich) and 20 μM Taxol (Sigma-Aldrich) or Docetaxel (Sanofi-Aventis) in MRB80 buffer (pH 6.8, 80 mM K-PIPES, 1 mM EGTA and 4 mM MgCl_2_) at 37°C for 30 minutes. After polymerization, GTP-tubulin-Taxol mix was diluted 5 times with prewarmed 20 μM Taxol or Docetaxel made in MRB80 buffer and centrifuged at 16200xg for 15 min at room temperature. The microtubule pellet was resuspended in prewarmed 20 µM Taxol solution in MRB80 buffer and stored at room temperature in the dark covered with aluminium foil.

#### In vitro reconstitution of microtubule nucleation

Flow chamber was assembled using plasma-cleaned glass coverslip and microscopic slide attached together using double-sided tape. These chambers were then functionalized by 5 min incubation with 0.2 mg/ml PLL-PEG-biotin (Susos AG, Switzerland) followed by 5 min incubation with 1 mg/ml NeutrAvidin (Invitrogen) in MRB80 buffer. Next, biotinylated-anti-GFP nanobody was attached to the coverslip through biotin-NeutrAvidin links by incubating the chamber for 5 min. During this incubation, γ-TuRC-GFP was diluted 10 times and either preincubated with nucleation-promoting factors or MRB80 buffer for 3 min and then immobilized on the GFP-nanobody coated coverslips by incubating it with flow chamber by 3 min. Non-immobilized γ-TuRC was washed away with MRB80 buffer and flow chambers were further incubated with 0.8 mg/ml k-casein to prevent non-specific protein binding. The nucleation mix with or without proteins (MRB80 buffer supplemented with 17.5 μM or 25 μM porcine brain tubulin, 50 mM or 80 mM (for CAMSAP assays) KCl, 1mM GTP, 0.5 mg/ml k-casein, 0.1% methylcellulose, and oxygen scavenger mix (50 mM glucose, 400 mg/ml glucose-oxidase, 200 mg/ml catalase, and 4 mM DTT)) were added to the flow chambers after centrifugation in an ultracentrifuge (Beckman Airfuge) at 119,000xg for 5 minutes. Concentrations of the proteins and composition of the nucleation mix have been indicated in either figure or figure legends. The flow chambers were then sealed with high-vacuum silicone grease (Dow Corning), and three consecutive 10 min time-lapse videos were acquired after 2 min incubation (time, t=0) of the flow chambers with the nucleation reactions on total internal reflection fluorescence microscope stage at 30 °C. All tubulin products were from Cytoskeleton.

#### Capping assays

For capping assays, 2 parallel flow chambers on the same coverslip were functionalized as mentioned above and then incubated with GMPCPP- and Taxol- or GMPCPP- and Docetaxel-stabilized microtubules for 2 min for their attachment to the coverslips through biotin-neutravidin links. Flow chambers were then incubated with 0.8 mg/ml k-casein to prevent non-specific protein binding and a master mix of reaction mix (5 μM tubulin, 50 mM KCl, 1mM GTP, 0.5 mg/ml k-casein, 0.1% methylcellulose, and oxygen scavenger mix (50 mM glucose, 400 mg/ml glucose-oxidase, 200 mg/ml catalase, and 4 mM DTT) in MRB80 buffer centrifuged in an Airfuge for 5 min at 119,000xg) supplemented with γ-TuRC (GCP3-GFP) and with or without 30 nM mCherry-CDK5RAP2 was prepared and divided into 2 for adding it to the parallel chambers. The flow chambers were then sealed with high-vacuum silicone grease and three 5 min time-lapse videos were acquired from both the parallel chambers alternatively after 2 min incubation (time, t=0) on TIRF microscope stage at 30 °C. Time interval was kept 20 s.

### Image acquisition, processing and data analysis

#### Widefield microscopy

Fixed HEK293T cells were imaged on a Nikon Eclipse Ni upright widefield fluorescence microscope equipped with a Nikon DS-Qi2 camera (Nikon), an Intensilight CHGFI precentered fiber illuminator (Nikon), ET-DAPI and ET-EGFP filters (Chroma), controlled by Nikon NIS Br software. Slides were imageg using a Plan Apo Lambda 60x NA 1.4 oil objective (Nikon).

#### TIRF microscopy

In vitro assays were imaged on an iLas2 TIRF microscope setup as described previously^56^. Briefly, ILas2 system (Roper Scientific, Evry, France) is a dual laser illuminator for azimuthal spinning TIRF illumination and powered with a custom modification for targeted photomanipulation. This system was installed on the Nikon Eclipse Ti-E inverted microscope with the perfect focus system. This microscope was equipped with Nikon Apo TIRF 100x 1.49 N.A. oil objective (Nikon), CCD camera CoolSNAP MYO M-USB-14-AC (Roper Scientific), EMCCD Evolve mono FW DELTA 512×512 camera (Roper Scientific) with the intermediate lens 2.5X (Nikon C mount adaptor 2.5X), 150 mW 488 nm laser, 100 mW 561 nm laser and 49002 and 49008 Chroma filter sets and controlled with MetaMorph 7.10.2.240 software (Molecular Devices). The final magnification using Evolve EMCCD camera was 0.064 μm/pixel and for CoolSNAP Myo CCD camera it was 0.045 μm/pixel. Temperature was maintained at 30 °C to image the in vitro assays using a stage top incubator model INUBG2E-ZILCS (Tokai Hit). Time-lapse movies were acquired using a CoolSNAP Myo CCD camera (Roper Scientific) at either 5 s (for nucleation assays without CAMSAPs) or 20 s (for assays with CAMSAPs) time interval with 200 ms, 300 ms and 500 ms exposure time for 642 nm, 561 nm and 488 nm illumination respectively for 10 minutes, while time-lapse images acquired on more sensitive Photometrics Evolve 512 EMCCD camera (Roper Scientific) were taken at 3 s time interval with 100 ms exposure time for 10 minutes.

#### γ-TuRC nucleation efficiency

For each independent nucleation assay, γ-TuRCs that nucleated microtubules or were anchored to an already nucleated microtubule within 10 min of a time-lapse movie were manually counted. Also, all the γ-TuRC (GCP3-GFP or GCP6-GFP) particles were detected and counted from the first or second frame of that time-lapse movie using an open-source ImageJ plugin ComDet v.0.5.4 https://github.com/ekatrukha/ComDet and the percentage of active ones out of the total was calculated and plotted for nucleation efficiency. From each independent assay, nucleation efficiency was calculated for three consecutive 10 min time-lapse movies from three different fields of view and plotted as 0-10, 10-20 and 20-30 min.

#### Microtubule growth dynamics analysis

Images and movies were processed using Fiji (https://imagej.net/Fiji). Kymographs from the in vitro reconstitution assays were generated using the ImageJ plugin KymoResliceWide v.0.4 https://github.com/ekatrukha/KymoResliceWide. Microtubule dynamics parameters viz. plus-end growth rate and catastrophe frequency were determined from kymographs using an optimized version of the custom made JAVA plugin for ImageJ as described previously^57, 62^. The relative standard error for catastrophe frequency was calculated as described in^56^.

#### Single-molecule intensity analysis of surface attached γ-TuRC

To estimate the number of GCP3-GFP molecules in γ-TuRC, three parallel flow chambers were assembled on the same plasma-cleaned glass coverslip. The three chambers were incubated with the dilutions of GFP protein (monomeric), GFP-EB3 (dimeric) and GCP3-GFP (test) strongly diluted to single molecules level. Flow chambers were then washed with MRB80 buffer, sealed with vacuum grease and immediately imaged using TIRF microscope. Samples were focussed first in one area and 15-20 images of unexposed coverslip regions were acquired with 100 ms exposure time. Acquisition settings were kept constant for the three parallel chambers. Single molecule fluorescence puncta were detected, measured and fitted with 2D Gaussian function using custom written ImageJ plugin DoM_Utrecht v.1.1.6 (https://github.com/ekatrukha/DoM_Utrecht). The fitted peak intensity values were used to build fluorescence intensity histograms. The histograms were fitted to Gaussian distributions using GraphPad Prism 9.

#### Single-molecule intensity analysis of γ-TuRC from CAMSAP3 assays

All the γ-TuRC (GCP3-GFP) particles immobilized on coverslip using biotinylated-GFP-nanobody within a field of view were detected from the first frame or second frame of a 10 min time-lapse movie using above mentioned ImageJ plugin ComDet. Integrated intensity values for individual active γ-TuRCs that nucleated microtubules and either colocalized or not colocalized with CAMSAP3 or dissociated from microtubules after CAMSAP3 recruitment were manually extracted and normalized to the average integrated intensity of total active γ-TuRCs that nucleated microtubules within that field of view.

#### Colocalization frequency of γ-TuRC and nucleation-promoting factors

All the γ-TuRC (GCP3-GFP) particles immobilized on coverslip within a field of view were detected from the first frame or second frame of 10 min time-lapse movies or single images (for assays without tubulin) using above mentioned ImageJ plugin ComDet and their colocalization percentage with the respective nucleation-promoting factor was calculated using the colocalization function of this ComDet plugin.

#### Ananlysis of γ-TuRC-CAMSAPs colocalization and microtubule release frequency

All the active γ-TuRCs that nucleated or anchored a microtubule were counted manually. Each of these γ-TuRC-anchored microtubule minus-ends were monitored for any binding of CAMSAPs within 10 min of a time-lapse movie and were scored manually if recruited CAMSAPs and plotted. Out of these CAMSAP-colocalized γ-TuRC-anchored minus-ends, the minus ends that started growing during this 10 min duration were quantified as released microtubules. For quantification of % of micrtotubules released from active γ-TuRC, all the γ-TuRC-anchored minus ends that started growing within 10 min were counted as released microtubules.

#### γ-TuRC capping efficiency

For quantification of γ-TuRC capping efficiency, stabilized microtubules were monitored for 5 min during a time-lapse movie and if one of the ends of a microtubule was stably bound to γ-TuRC for at least 2 min, that microtubule was scored as γ-TuRC-capped microtubule.

### Statistical analysis

All statistical details of experiments including the definitions, exact values of number of measurements, precision measures and statistical tests performed are mentioned in the figure legends. All experiments were repeated at least three times. Data processing and statistical analysis were done in Excel and GraphPad Prism 9 (GraphPad Software). Significance was defined as: ns =not significant, *p < 0.05, **p < 0.01, ***p < 0.001 and ****p < 0.0001.

## Legends to Supplementary Figures and Videos

**Extended data Fig. 1, related to Figure 1.**
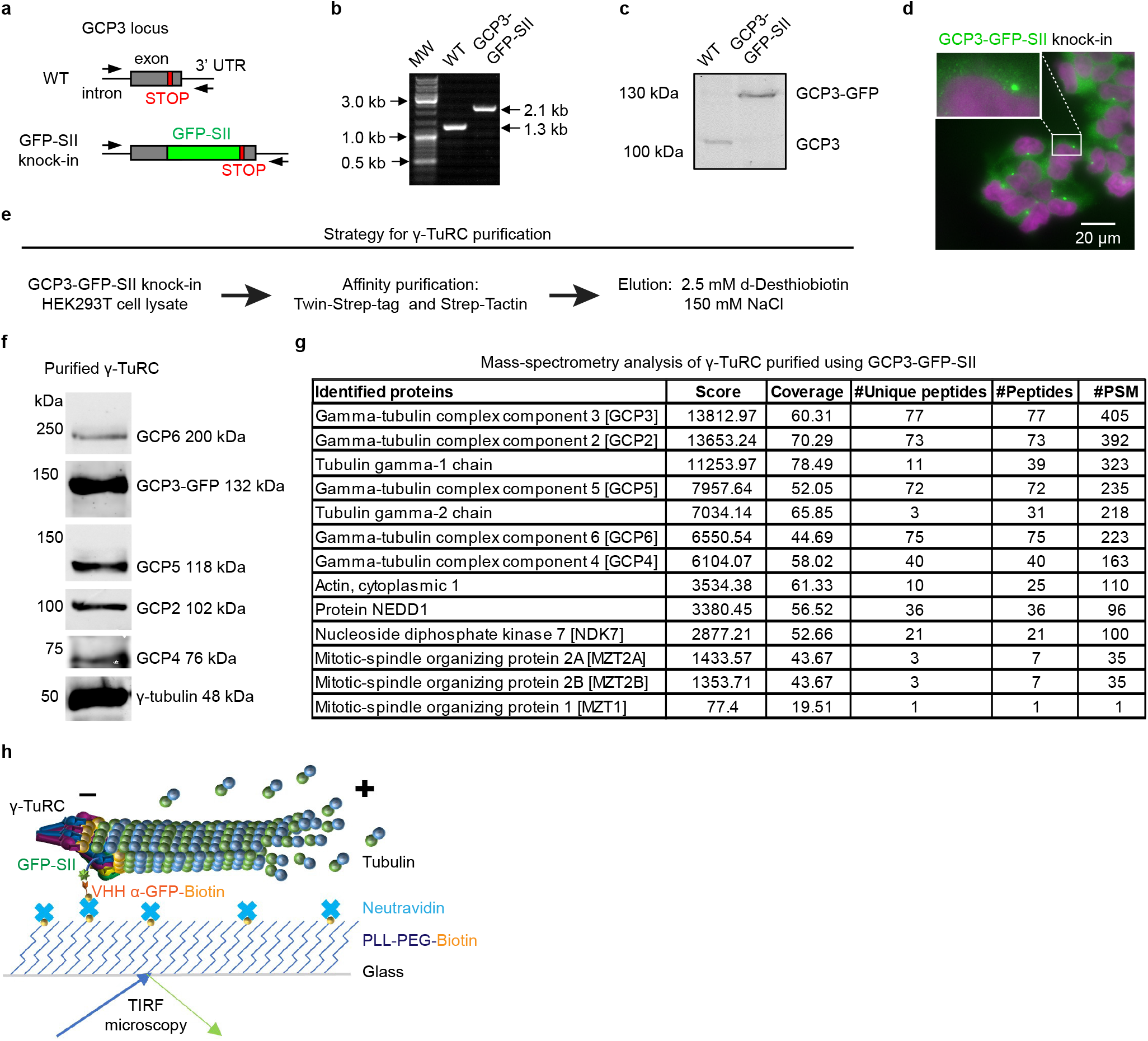
Characterization of HEK293T GCP3-GFP-SII homozygous knock-in cell line and γ-TuRC purified using GCP3-GFP-SII. **a,** Schematic showing GCP3 gene locus and knock-in strategy. **b,** 1% Agarose gel showing genomic-DNA PCR products for wild type HEK293T cells and GCP3-GFP-SII knock-in cells. MW, molecular weight DNA ladder; WT, wild type. **c,** Western blot for wild type and GCP3-GFP-SII knock-in HEK293T cell lysate, blotted using mouse anti-GCP3 antibody. **d,** Widefield fluorescent image of fixed HEK293T GCP3-GFP-SII homozygous knock-in cells showing GFP fluorescence (green). Nuclei (magenta) were stained with DAPI. **e,** Strategy for γ-TuRC purification. **f,** Western blot results showing the presence of all the core components in our γ-TuRC purified using GCP3-GFP-SII using antibodies against GCP6, GFP, GCP5, GCP2, GCP4 and γ-tubulin. **g,** Mass spectrometry results showing the presence of all the core components in γ-TuRC purified using GCP3-GFP-SII. **h,** Schematic showing experimental TIRF microscopy setup for in vitro reconstitution of microtubule nucleation from γ-TuRC.

**Extended data Fig. 2, related to Figure 1.**
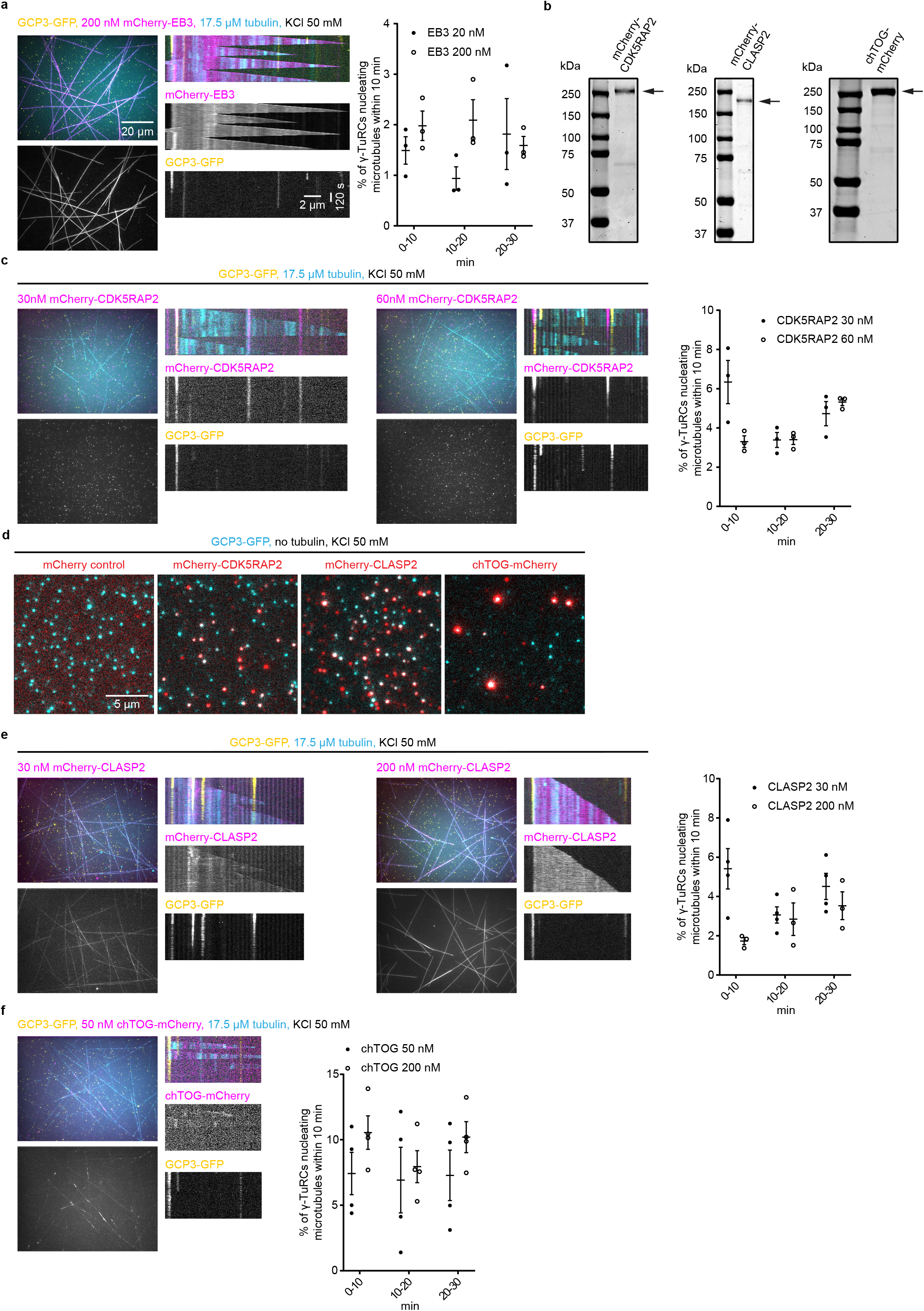
Characterization of effective concentrations of nucleation-promoting factors. a,c,e,f, Left: maximum intensity projections and representative kymographs illustrating microtubule dynamics in 10 min time-lapse videos, acquired after 20 min of incubation, showing microtubules (cyan) nucleated from γ-TuRC (GCP3-GFP, yellow) in the presence of either 17.5 μM tubulin (17 μM unlabeled porcine tubulin and 0.5 μM HiLyte647-tubulin), 50 mM KCl and together with indicated concentrations of indicated proteins (magenta) and without any preincubation: 200 nM mCherry-EB3 (**a**); or 30 nM and 60 nM mCherry-CDK5RAP2 (**c**); or 30 nM and 200 nM mCherry-CLASP2 (**e**); or 50 nM chTOG-mCherry (**f**). Right: Quantification of average microtubule nucleation efficiency of γ-TuRC as indicated: 200 nM mCherry-EB3 (n=3); 30 nM (n=3) or 60 nM (n=3) mCherry-CDK5RAP2; 30 nM (n=4) or 200 nM mCherry-CLASP2 (n=3); 50 nM (n=4) or 200 nM chTOG-mCherry (n=4); where n is the number of independent experiments analyzed, also see Fig. 1c-h. The plots present mean ± s.e.m., and each data point represents a single field of view from which % of γ-TuRCs nucleating microtubules were quantified for the given time point from an individual experiment. Data points at 0-10 min were acquired from a smaller field of view and cannot be directly compared to the data points at 10-20 min and 20-30 min acquired from a bigger field of view (shown in the panels on the left). Data points for concentrations already shown in Fig. 1c,e-h, have been taken from Fig. 1d and replotted here for direct comparison. **b,** Coomassie-stained SDS-PAGE gels loaded with purified mCherry-CDK5RAP2 or mCherry-CLASP2 or chTOG-mCherry. **d,** Representative images showing colocalization of γ-TuRC (cyan, GCP3-GFP) with indicated proteins (red) in the presence of 50 mM KCl and no soluble tubulin.

**Extended data Fig. 3, related to Figure 2.**
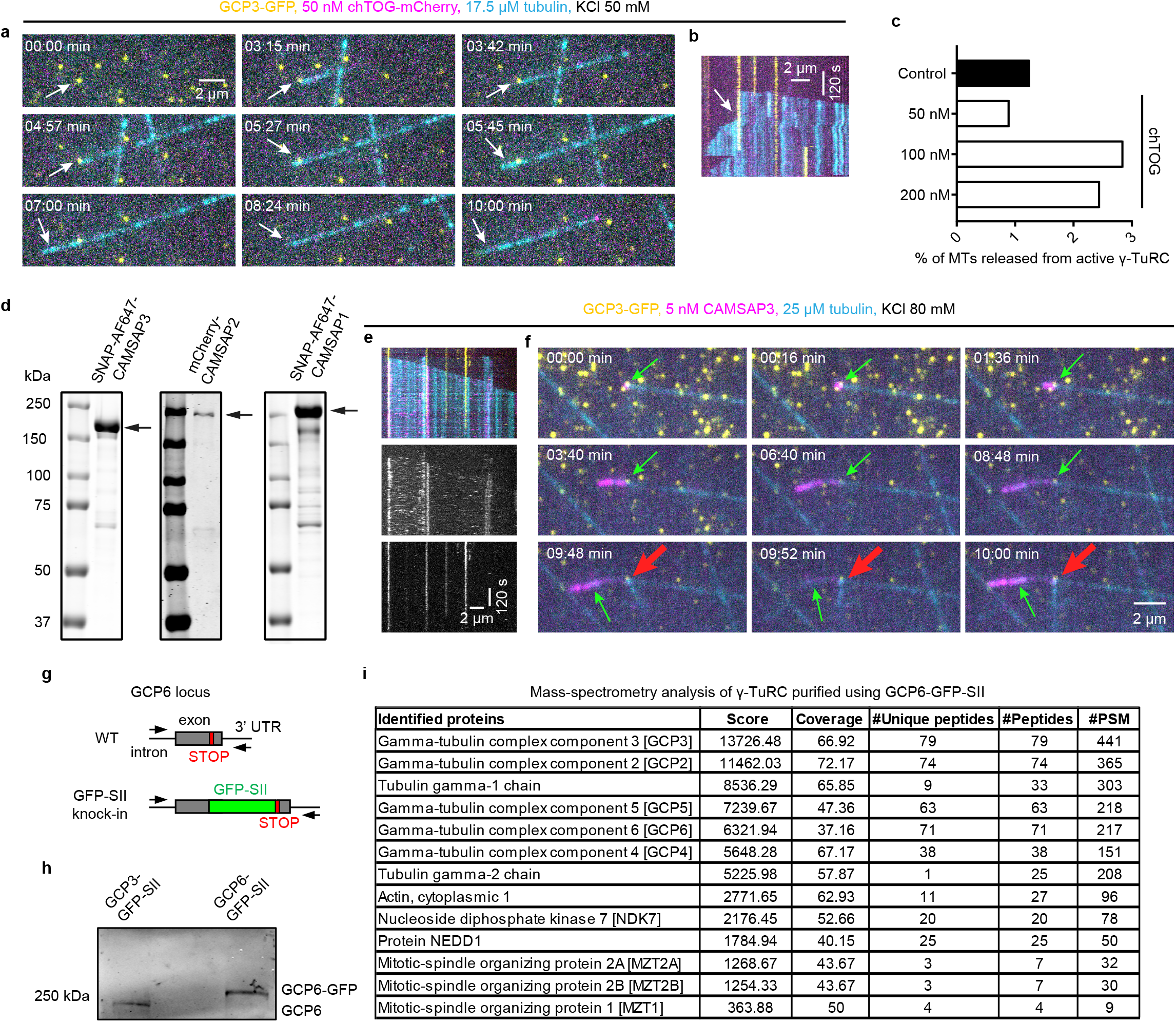
Characterization of purified CAMSAPs and GCP6-tagged γ-TuRC and microtubule release from γ-TuRC. **a,b,** Still frames (at indicated time points in min) (**a**) and representative kymograph (**b**) from a 10 min time-lapse video showing microtubule (cyan) nucleation (magenta) and subsequent microtubule release from γ-TuRC (GCP3-GFP, yellow) in the presence of 50 nM mCherry-chTOG (magenta), 17.5 μM tubulin (17 μM unlabeled porcine tubulin and 0.5 μM HiLyte647-tubulin). Thin arrows indicate microtubule minus-end. **c,** Plot showing frequency of microtubule release from active γ-TuRC over 10 min duration in the presence of either 17.5 μM tubulin alone (control, n=52, pooled from eight independent experiments); or together with 50 nM (n=280, pooled from four independent experiments); or 100 nM (n=183, pooled from four independent experiments); or 200 nM mCherry-chTOG (n=301, pooled from four independent experiments); where n is the number of active γ-TuRCs analyzed. Representative images are shown in **a**. **d,** Coomassie-stained SDS-PAGE gels loaded with purified SNAP-AF647-CAMSAP3, mCherry-CAMSAP2 or SNAP-AF647-CAMSAP1. **e,f,** Two different examples: kymographs (**e**) and still frames (at indicated time points in min) from a 10 min time-lapse video showing γ-TuRC-CAMSAP3 interplay at the γ-TuRC-anchored microtubule minus-ends under indicated experimental conditions. Example 1 (kymographs) illustrates occasions when CAMSAP3 (magenta) fails to displace γ-TuRC (GCP3-GFP, yellow) from γ-TuRC-anchored microtubule (cyan) minus-end that recruited CAMSAP3. Example 2 (still frames) illustrates microtubule re-nucleation (thick red arrows) from the same γ-TuRC that released previously nucleated microtubule (thin green arrows) upon CAMSAP3 binding and minus-end growth. Experimental conditions are same as shown in Fig. 2e, i.e., 25 μM tubulin (24.5 μM unlabeled porcine tubulin and 0.5 μM rhodamine-tubulin), 80 mM KCl and 5 nM SNAP-AF647-CAMSAP3. **g,** Schematic showing GCP6 gene locus and knock-in strategy. **h,** Western blot for GCP3-GFP-SII and GCP6-GFP-SII knock-in HEK293T cell lysate, blotted using mouse anti-GCP6 antibody. **i,** Mass-spectrometry results showing the presence of all the core components in our γ-TuRC purified using GCP6-GFP-SII.

**Extended data Fig. 4, related to Figure 3.**
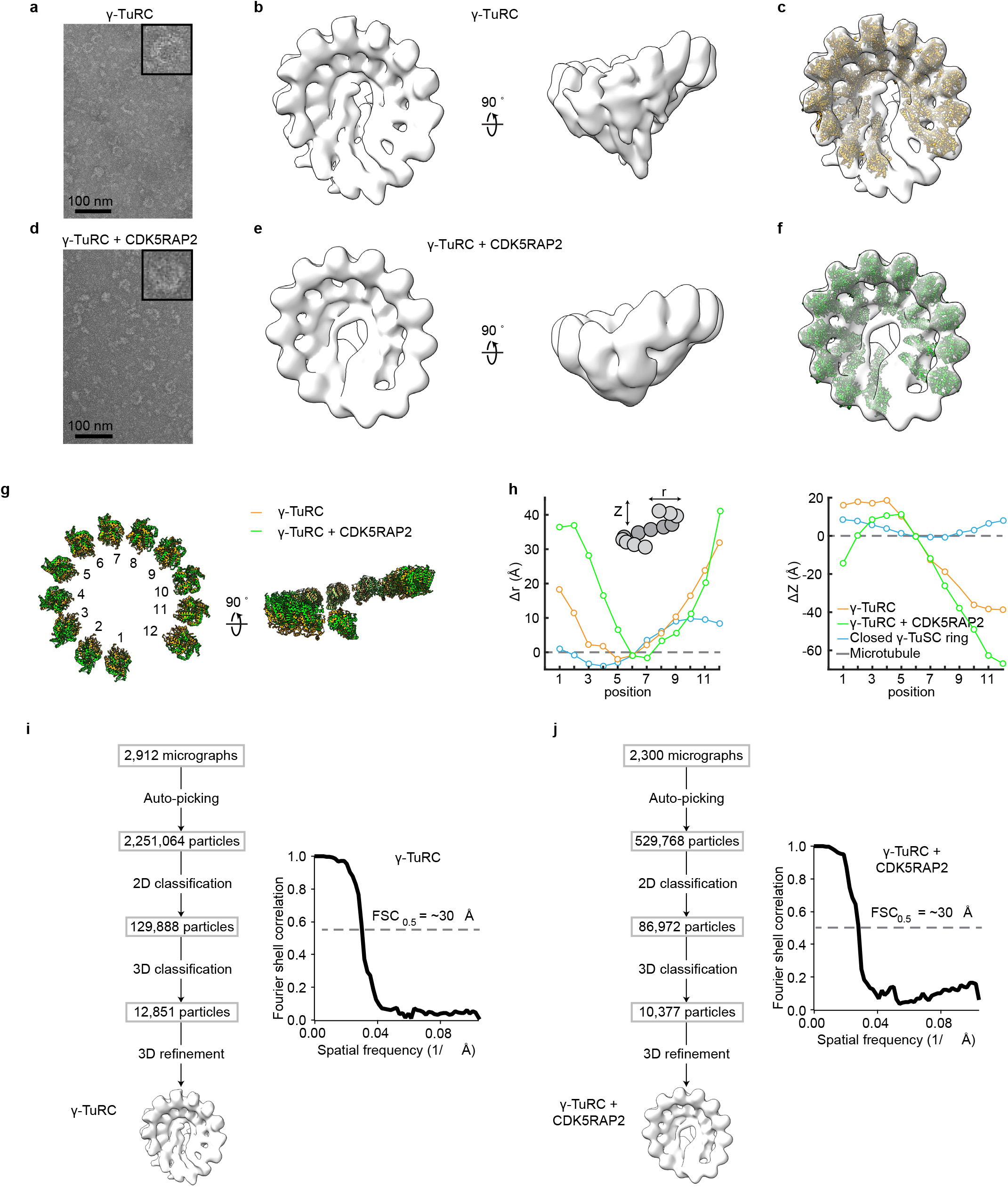
EM-based characterization of γ-TuRC in the absence or presence of CDK5RAP2. **a,** Transmission EM (TEM) micrograph of negatively stained γ-TuRC. Inset shows a 4X magnified view of a single γ-TuRC. **b,** Two views of a 3D reconstruction of the γ-TuRC from negative-stain EM data. **c,** Rigid body fit of repeating γ-tubulin/GCP2 subcomplexes (from PDB ID: 6V6S^20^) individually docked into the γ-TuRC density map. Fits for two subcomplexes at the γ-TuRC “seam” were not reliable; and are therefore, omitted for clarity. **d,** Transmission EM (TEM) micrograph of negatively stained γ-TuRC prepared in complex with 120 nM CDK5RAP2. Inset shows a 4X magnified view of a single γ-TuRC. **e,** Two views of a 3D reconstruction of the γ-TuRC + CDK5RAP2 preparation from negative-stain EM data. **f,** Rigid body fit of repeating γ-tubulin/GCP2 subcomplexes (from PDB ID: 6V6S^20^) individually docked into the γ-TuRC + CDK5RAP2 density map. As in **c**, fits for two subcomplexes at the γ-TuRC “seam” were not reliable; and are therefore, omitted for clarity. **g,** Two views of the γ-tubulin rings from rigid body fitted models in **c** (γ-TuRC) and **f** (γ-TuRC + CDK5RAP2). **h,** Plots of the change in helical radius (*r)* and helical pitch (*Z*) relative to *β*-tubulin in the 13-protofilament microtubule lattice (grey dashed line; PDB ID: 2HXF ^43^and EMD-5193 ^44^) calculated for γ-tubulin rings from γ-TuRC alone (orange), γ-TuRC + 120 nM CDK5RAP2 (green), and the γ-TuSC oligomer in the “closed” state (blue; PDB ID: 5FLZ^42^). See Methods for analysis details. **i,** Left: Processing workflow for generating a negative-stain EM 3D reconstruction of γ-TuRC. Right: Unmasked FSC curve for the γ-TuRC reconstruction. FSC =0.5 is indicated by a dashed gray line, and an estimate of the corresponding resolution is indicated. **j,** Left: Processing workflow for generating a negative-stain EM 3D reconstruction of γ-TuRC in the presence of 120 nM CDK5RAP2. Right: Unmasked FSC curve for the γ-TuRC + CDK5RAP2 reconstruction. FSC=0.5 is indicated by a dashed gray line, and an estimate of the corresponding resolution is indicated.

## Supplementary Video Legends

**Supplementary Video 1. CAMSAP3 mediated microtubule release from γ-TuRC.** 10 min time-lapse videos illustrating CAMSAP3 binding to γ-TuRC-anchored (left & middle: GCP3-GFP; right: GCP6-GFP) minus ends and subsequent elongation of the CAMSAP3-stabilized minus ends in the presence of 25 μM tubulin (24.5 μM unlabeled porcine tubulin and 0.5 μM rhodamine-tubulin), 80 mM KCl and 5 nM SNAP-AF647-CAMSAP3. Time-lapse images were acquired using TIRF microscope at 20 s (left, constituting 31 frames) and 3 s (middle and right, constituting 201 frames) time intervals.

**Supplementary Video 2. γ-TuRC displacement from microtubule minus ends by CAMSAP3 in the presence of nucleation-promoting factors.** 10 min time-lapse videos illustrating CAMSAP3 binding to γ-TuRC-anchored minus ends and subsequent elongation of the CAMSAP3-stabilized minus ends in the presence of 25 μM tubulin (24.5 μM unlabeled porcine tubulin and 0.5 μM rhodamine-tubulin), 80 mM KCl, 5 nM SNAP-AF647-CAMSAP3 and indicated nucleation-promoting factors (30 nM mCherry-CDK5RAP2 or 30 nM mCherry-CLASP2 or 200 nM chTOG-mCherry). Time-lapse images were acquired using TIRF microscope at 20 s time interval constituting 31 frames.

**Supplementary Video 3. γ-TuRC detachment from microtubule minus ends by CAMSAP2 or CAMSAP1.** 10 min time-lapse videos illustrating CAMSAPs binding to γ-TuRC-anchored minus ends and subsequent elongation of the CAMSAPs-stabilized minus ends in the presence of 25 μM tubulin (∼2% labeling), 80 mM KCl and 40 nM mCherry-CAMSAP2 or 5 nM SNAP-AF647-CAMSAP1. Time-lapse images were acquired using TIRF microscope at 20 s time interval constituting 31 frames.

## Supplementary Table Legends

**Supplementary Table 1. Mass-spectrometry results of purified GCP3-GFP-SII.** List of proteins that were co-purified with endogenously tagged GCP3-GFP-SII contains all the core components of γ-TuRC, identified by mass spectrometry.

**Supplementary Table 2. Mass-spectrometry results of purified CDK5RAP2.** List of proteins identified by mass spectrometry that were co-purified with SII-mCherry-CDK5RAP2.

**Supplementary Table 3. Mass-spectrometry results of purified CLASP2.** List of proteins identified by mass spectrometry that were co-purified with SII-mCherry-CLASP2.

**Supplementary Table 4. Mass-spectrometry results of purified chTOG.** List of proteins identified by mass spectrometry that were co-purified with chTOG-mCherry-SII.

**Supplementary Table 5. Mass-spectrometry results of purified GCP6-GFP-SII.** List of proteins that were co-purified with endogenously tagged GCP6-GFP-SII contains all the core components of γ-TuRC, identified by mass spectrometry.

**Source data 1.**
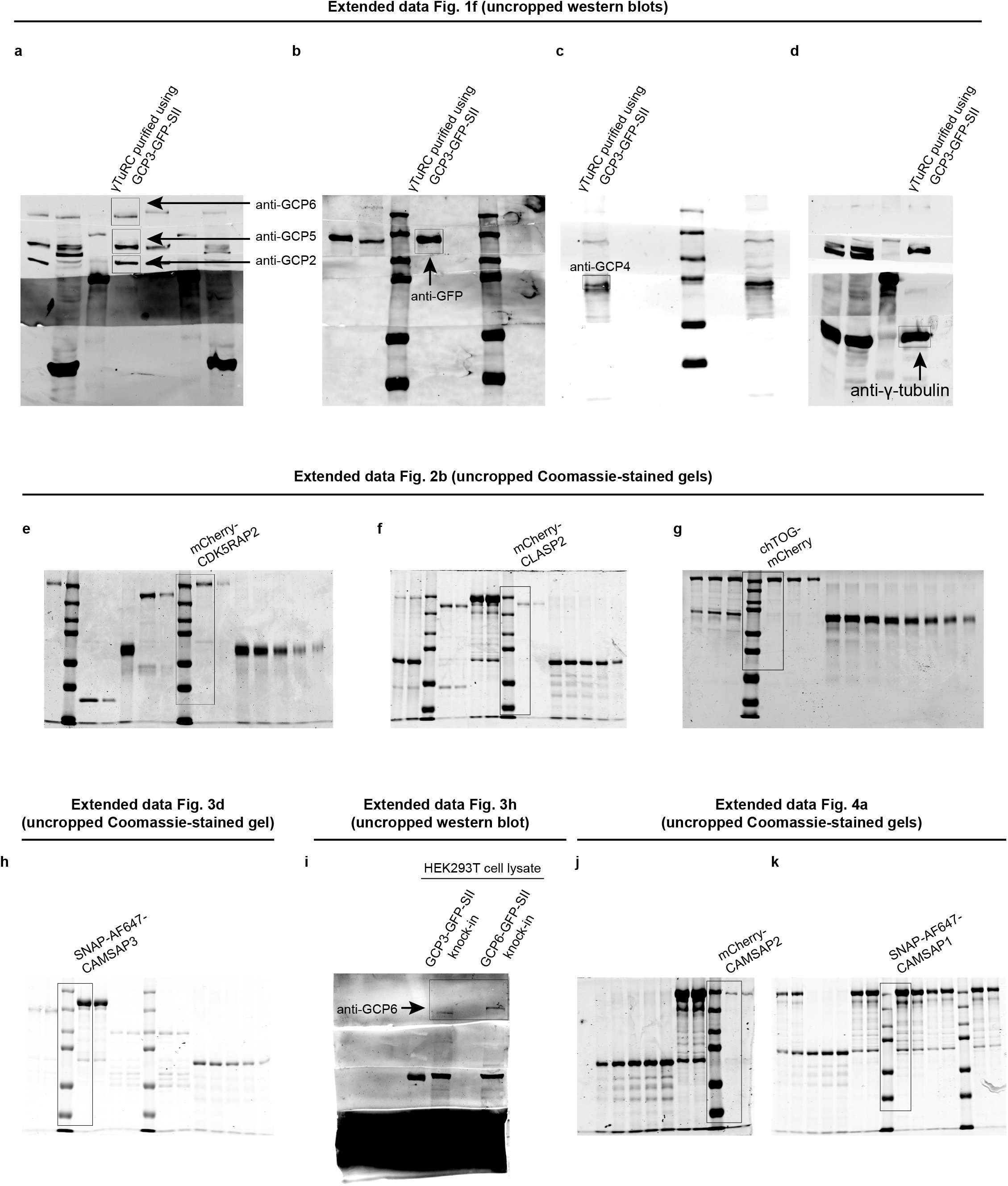

